# High-Throughput Machine Learning-Aided Antibody Discovery for Cell Surface Antigens

**DOI:** 10.1101/2025.05.15.650607

**Authors:** Deepash Kothiwal, Aaron W. Kollasch, Nicholas Hollmer, Anita Ghosh, Roushu Zhang, Murali Anuganti, Steffanie B. Paul, Yvrick Zagar, Mina Abdollahi, Zachary Anderson, Filmawit Belay, Matt Salotto, Sophia Ulmer, Youssef Atef Abdelalim, Satyendra Kumar, Mahesh Vangala, Chang Yang, Alain Chedotal, Joseph G. Jardine, Andre A. R. Teixeira, Deborah J. Moshinsky, Haisun Zhu, Shaotong Zhu, Timothy A. Springer, Debora S. Marks, Rob Meijers

**Affiliations:** Institute for Protein Innovation, Boston, MA 02115, USA; Department of Systems Biology, Harvard Medical School, Boston, MA, USA; Sorbonne Université, INSERM, CNRS, Institut de la Vision, Paris, France; Institut de Pathologie, Groupe Hospitalier Est, Hospices Civils de Lyon, Lyon, France; MeLiS, CNRS UMR5284, Inserm U1314, University Claude Bernard Lyon 1, Lyon, France; Department of Immunology and Microbiology, Scripps Research Institute, La Jolla, CA 92037, USA; IAVI Neutralizing Antibody Center, Scripps Research Institute, La Jolla, CA 92037, USA; Program in Cellular and Molecular Medicine, Boston Children’s Hospital, Boston, MA 02115, USA; Department of Biological Chemistry and Molecular Pharmacology, Harvard Medical School, Boston, MA 02115, USA; Broad Institute of Harvard and MIT, Cambridge, MA, USA

## Abstract

Machine learning (ML) has the potential to revolutionize antibody design and selection, but its success depends on access to extensive, well-curated datasets of antibody-antigen interactions. To address this need, we developed a synthetic Fab yeast display library optimized for seamless ML integration, focusing on sequence diversity within the CDRH3 loop. The library incorporates key sequence features derived from human B cell repertoires essential for efficient antibody generation captured in a compact antigen recognition module (ARM) format. Built using the VH1-69 heavy chain and four light chains, the library was evaluated against ten human and murine cell surface antigens, including PD-L1, TIGIT, and ROBO1. This approach yielded hundreds of antibodies with robust biophysical properties, validated for functional performance in flow cytometry and immunohistochemistry. Furthermore, ML analysis identified additional antibodies for ROBO2 and PD-L2 from the aggregate sequencing data, demonstrating utility for hybrid *in silico* and experimental workflows. We provide a publicly accessible dataset comprising more than 68,000 Fab sequences and 486 characterized antibodies. This study establishes an ML-compatible framework designed to accelerate and streamline antibody discovery and development.

## Introduction

Antibody display technologies provide promising alternatives for the discovery of antibodies with properties that rival monoclonal antibodies derived from patients or animal immunization^1,2^. The diversity of the antibody repertoire of the human immune system is recreated in a synthetic controlled framework using singular biological entities such as yeast cells or phage that can be manipulated with molecular biology techniques to display a large variety of binders for exposure to antigen^3^. A purely synthetic display library offers the advantage of a robust and controlled antibody discovery process, which is ideal for developing machine learning algorithms to assist in hybrid in silico/experimental antibody discovery^4^. In contrast to species-specific immunization, synthetic libraries offer access to highly evolutionarily conserved antigens, e.g. in the nervous system and in development, that are currently underserved by traditional monoclonal antibodies. However, state-of-the-art display libraries for therapeutic antibody development often rely on patient or animal-derived complementary determining regions (CDRs) of high complexity, using a combination of selection techniques that complicate the development of datasets for large language models^2^.

It has been shown that the heavy chain CDRH3 loop provides unique and specific antigen recognition *in vivo*^5^, forming the core module of antigen recognition^6^. Reducing the antibody paratope sequence space from all six CDRs to the CDRH3 alone significantly constrains antigen/antibody interaction language models and accelerates *in silico* antibody development. Synthetic libraries that only explore diversity in the CDRH3 region have been produced previously^7–11^, using trinucleotide mutagenesis technology with one fixed amino acid frequency for all CDRH3 oligos^12^. The use of fixed, random sequence repertoires is problematic, because it introduces amino acid motifs that cause polyreactivity, aggregation and degradation that are removed from a natural immune repertoire through B cell selection. Combinatorial variant library synthesis methods allow the editing of selective sequence repertoire features with high diversity into synthetic antibody libraries, enhancing their biochemical properties^13,14^.

Here, we describe a synthetic Fab yeast display library that incorporates position-specific amino acid frequencies for the CDRH3 region derived from a naive B cell repertoire, with additional removal of amino acid motifs that could cause antibody aggregation and polyreactivity^2^. The CDRH3 sequence is followed by a nucleotide-level barcode to track light chain pairing, which together represent the antigen recognition module (ARM) of the antibody. The ARM is encoded by a compact nucleotide sequence amenable to deep sequencing, thus providing a convenient shorthand of the antibody paratope. The Fab library was screened against ten biologically significant and therapeutically relevant cell surface targets, yielding antibodies with favorable biophysical and binding properties for all ten. We capitalize on the extensive NGS data set of antibody candidates obtained for PDL2 and ROBO2 to develop a machine learning protocol that identifies low-frequency Fab clones with favorable binding properties. These results highlight the potential of our minimal Fab library approach to support hybrid in silico/experimental antibody discovery.

## Results

### Design and sequence analysis of the antigen recognition module (ARM)

The ARM library was designed as a short nucleotide sequence (<100 nucleotides) covering the CDRH3 region and the adjacent framework region 4, which contained a nucleotide barcode to track heavy/light chain pairing (Fig. 1A). Oligo pools were designed separately for each CDRH3 length, which ranged from 11 to 17. Each position had a unique amino acid frequency derived from 9.5 million unique CDRH3 sequences from naive B cells obtained from the OAS database^15^ (Supplementary Fig. S1A). Diversity in amino acid frequency was applied to IMTG positions 107 to 114, with additional sampling of highly frequent amino acids in positions 115 and 117 (Fig. 1A, 1B and Supplementary Fig. S1B). To compensate for potential over- or undersampling of the CDRH3 loops, we adjusted the fractions for each length (Fig. 1B). In addition, cysteines, methionines and amino acid motifs with undesired biophysical and chemical properties were excluded from the oligo pool (Supplementary Table S1).

**Figure 1.**
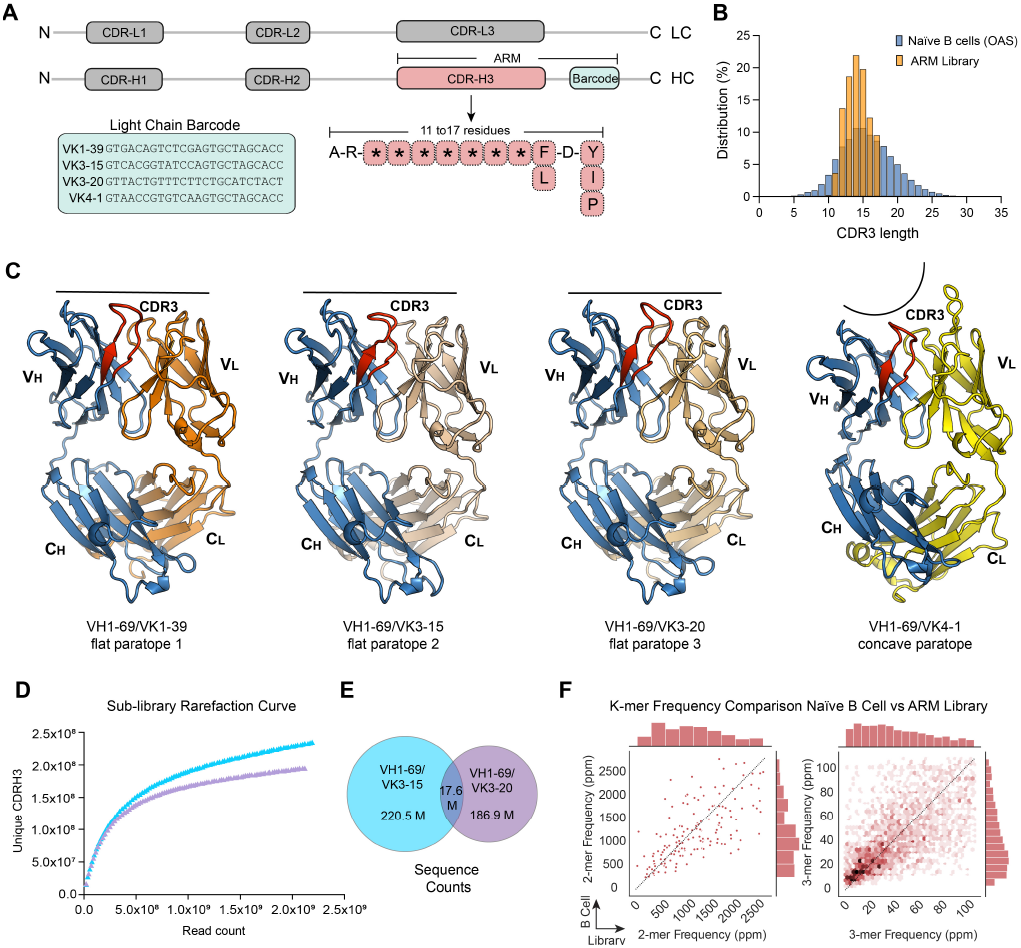
Construction of the ARM library for Fab yeast display. **(A)** Antigen recognition module (ARM) schematic covering the hypervariable CDRH3 region of the heavy chain (HC) followed by a light chain (LC) barcode that uses degenerate codons to insert silent mutations corresponding to the co-expressed light chain. **(B)** Histogram of the native B cell CDRH3 length distribution as obtained from the OAS database (blue) and CDRH3 length distribution of the VH1-69/VK3-15 ARM sublibrary (orange). **(C)** Alphafold3^38^ structural models of the Fab fragments. The CDRH3 region is colored red and a black bar emphasizes the shape of the antigen recognition surface as flat or concave. **(D)** Diversity accumulation plot for two sublibraries performed with Novaseq, displaying the number of unique CDRH3s versus the number of read counts. **(E)** Venn diagram of the sequence overlap between the two sublibraries. **(F)** K-mer frequency correlation between the naive B cell CDRH3 repertoire and the ARM library, expressed in parts per million for each k-mer for the VH1-69/VK3-15 sublibrary.

The library was constructed using the VH1-69 heavy chain germline sequence as a molecular scaffold. VH1-69 is a common antibody heavy chain germline that is well represented in robust therapeutic antibodies^2^ and broadly neutralizing antibodies for viral antigens^16^. The CDR1 and CDR2 sequence of the VH1-69 heavy chain were based on frequently occurring germline sequences observed in the OAS database (Supplementary Fig. S2A). Four VH1-69 heavy chain Fab sublibraries were created, each containing a unique barcode, to distinguish the paired light chain during the selection process (Fig. 1A and Supplementary Fig. S2B and S2C). Yeast populations for each sub-library were then mixed for antibody discovery, creating a repertoire of VH1-69/light chain pairs (Supplementary Fig. S2D). The VK1-39, VK3-15, VK3-20 light chains were chosen based on their prominence in therapeutic antibodies with favorable biophysical properties^2^ and present a relatively flat paratope (Fig. 1C). The VK4-1 light chain has a long CDRL1 loop and provides a more concave paratope, potentially enabling binding to alternative epitopes including peptides^17–19^.

Deep sequencing on two of the four VH1-69 sub-libraries revealed 390 million unique CDRH3 sequences. No dominant clones were observed, and most redundancies were limited to duplicates or triplicates. Due to incomplete oligo assembly, shorter 9mer and 10mer CDRH3 sequences were observed although they only contribute 0.3 % to the entire CDRH3 repertoire (Fig. 1B and Supplementary Table S2). Deleterious motifs in the CDRH3 region of the antigen recognition module were compared with a randomized set of CDRH3 sequences and the CDRH3 sequences derived from naïve B cells in the OAS database. The Fab library described here is largely devoid of deleterious motifs, whereas 60% of CDRH3 sequences in the random and naïve B cell libraries contained at least one deleterious motif (Supplementary Fig. S3A). Using rarefaction curves, the molecular diversity of each of these two VH1-69 sublibraries was estimated to be ∼2.5 x10^8^ unique clones (Fig. 1D). Sequence comparison of the CDRH3 pools from the two sublibraries revealed a relatively low overlap (9%), confirming that the parental oligo pool used for transformation in yeast possessed a far higher diversity (Fig.1E). This finding indicated that each sublibrary contributed independently to the molecular diversity, not only because each was paired with a different light chain, but also because the vast majority of CDRH3 sequences were unique. Since four sublibraries were generated, the true molecular diversity of the VH1-69 library was estimated to approach one billion unique Fabs.

Sequence analysis showed that the CDRH3 amino acid frequency distribution reflects that of the naïve B cell population reasonably well, except for minor variations due to the removal of liability motifs (Supplementary Fig. S1). The k-mer usage in the designed library correlated highly with k-mer usage in naïve B cell sequences with Spearman correlation values of 0.93, 0.78, 0.76 for 1-mer, 2-mers, 3-mers, respectively (Fig. 1F). The high correlation of 1-mers was expected, as the designed library was explicitly defined based on amino acid frequencies in naïve B cell sequences (Supplementary Fig. S3B). Some differences were observed between the designed and natural sequences. For example, in the 3-mer analysis, motifs containing glycine and/or tyrosine occurred much more frequently in naïve sequences than in the designed library, as a consequence of intentional removal of some of the Y and G-containing motifs during oligo design. In conclusion, the sequence constraints introduced through combinatorial oligo synthesis reproduced features of the amino acid frequencies observed in natural B cell libraries, while effectively reducing the occurrence of deleterious amino acid motifs that may cause polyreactivity, heterogeneity and aggregation of the antibodies (Supplementary Table S1).

### A high-throughput antibody discovery campaign delivered reagent & therapeutic grade antibodies for ten targets

To investigate the applicability of the minimalist Fab library for generating antibodies against a wide range of antigens, we carried out a single antibody discovery campaign against ten biomedically relevant cell surface targets in parallel. The targets included immune checkpoint inhibitors (PD-L1, PD-L2 and TIGIT)^20^, the lectin-like receptor LOX1^21^, a soluble Wnt signaling antagonist (Dickkopf-1)^22^, a cytokine receptor (IL23R)^23^, neuronal^24^ receptors (DCC, ROBO1 and ROBO2) and a human viral fusion-like protein (Syncytin-2)^25^. These targets were selected to challenge the library with diverse antigens that span high value biological targets (green ribbon cartoon portions, Fig. 2A) that varied in size (e.g., large: IL23R; small: TIGIT), sequence identity (e.g., high: ROBO1 and ROBO2, 94%; low: PD-L1 and PD-L2, 50%), and structural characteristics—including β-sandwich folds typical of immunoglobulin or fibronectin domains, mixed α/β domains (LOX1, DKK1, and Syncytin-2), and regions with substantial α-helical content (LOX1 and Syncytin-2) (Fig. 2A). Separate discovery campaigns were executed for a pool of the three flat paratope VH1-69/VK pairs and the VH1-69/VK4-1 concave paratope, to investigate if the overall paratope shape would affect Fab selection. Multiple rounds of magnetic and fluorescent cell sorting were performed with decreasing antigen concentrations to enrich Fab displaying yeast populations with strong binders (Fig. 2B & C, Supplementary Fig. S4 and S5).

**Figure 2.**
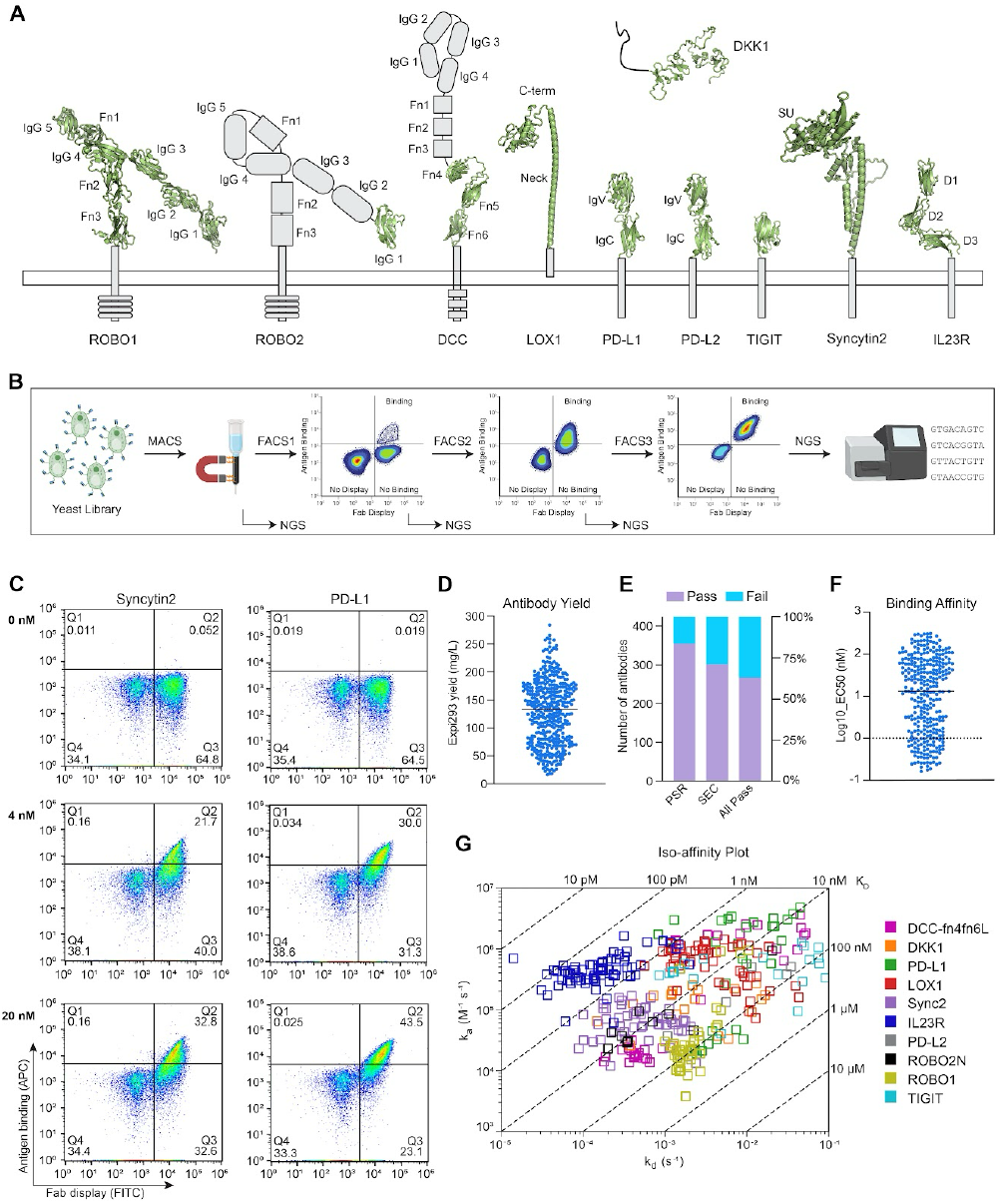
Antibody discovery campaign for ten glycoproteins. **(A)** Schematic of the ten glycoprotein antigens, showing the actual protein construct used in the discovery campaign as a ribbon in green. **(B)** Cell sorting scheme used to select yeast clones displaying Fabs specific for a particular antigen. The selection process involved one round of MACS followed by three rounds of FACS. After each FACS round, the selected yeast colonies were grown out and NGS sequenced, to track Fab enrichment during the selection process. (**C)** Examples of FACS plots for antigens Syncytin-2 and PD-L1 from the final FACS round, showing enrichment of yeast clones displaying intact Fabs responding to different concentrations of antigen. The display is measured on the x-axis through the presence of the myc epitope tag on the light chain of the Fab. The antigen binding response is measured on the y-axis through interaction with biotinylated antigen bound the strep-APC. **(D)** Antibody yields for all recombinant antibodies produced. **(E)** Pass/fail rates for the recombinant antibodies produced during affinity purification, in a polyspecificity reagent (PSR) assay and size exclusion chromatography (SEC). **(F)** Cell display antibody titration EC50 values for all generated antibodies. Iso-affinity plot showing k_a_, k_d_ and K_D_ derived from SPR measurements for all experimentally verified antibodies.

Yeast populations from each sorting step were deep sequenced by amplifying the ARM which included the CDRH3 region and the adjacent light chain bar code (Fig. 1A). We clustered CDRH3 sequences together that have the same length and shared more than 80% of the maximum BLOSUM62 score in pairwise sequence alignment. Based on CDRH3 clustering of the most frequent ARMs observed in the final FACS round, synthetic genes of the most prominent member of each selected CDRH3 cluster were ordered for 429 heavy chains. This method proved cost effective as each insert contained fewer than 100 nucleotides that were incorporated into an expression plasmid containing the remaining portion of the VH1-69 heavy chain. Human IgG1 antibodies were produced by coexpressing the VH1-69 synthetic genes with the corresponding light chain-containing plasmids, yielding 424 antibodies with sufficient production levels (Fig. 2D). The integrity of the antibodies was validated by size exclusion chromatography (SEC), and 302 antibodies met a stringent benchmark showing no aggregation or degradation (Fig. 2E & Supplementary Table S3). In parallel, polyreactivity was evaluated by ELISA, with 354 antibodies not showing reactivity to avidin, DNA, insulin or a membrane preparation derived from Sf9 insect cells and 285 antibodies passing both tests (Fig. 2E & Supplementary Table S3). Kinetics of the antibodies were measured using high-throughput surface plasmon resonance (SPR) and antibody potency was tested with a cell-based assay (Fig. 2F and 2G). These complementary techniques assessed binding to the original truncated antigen in SPR, and the complete ectodomain of the antigen in cell display. Many antibodies that passed the integrity and polyreactivity tests exhibited robust kinetics in SPR (103 antibodies with a K_D_ < 10 nM) and showed high potency in cell display (118 antibodies with EC_50_ < 25 nM) showing their promise as reagent antibodies and as early leads for therapeutics (Fig. 2F and 2G, Supplementary Fig. S6 and Supplementary Table S3).

Some of the best-performing antibodies from this campaign were tested in antibody reagent assays and compared to commercially available antibodies. Antibodies for TIGIT and LOX1, both important biomarkers monitored by flow cytometry, showed comparable potency to commercially available antibodies recommended for this application (Supplementary Fig. S7). ROBO1 and ROBO2 are axon guidance receptors and few good antibodies are available for immunohistochemistry experiments. While the antibodies were raised to human ROBO1 and ROBO2, cell display experiments showed strong cross-reactivity with murine versions of these receptors (Supplementary Fig. S6). We tested whether promising ROBO1 and ROBO2 antibodies could be used in immunohistochemistry antibody staining of mouse brain regions known to exhibit Robo1 and Robo2 expression. First, we converted the human IgG1 ROBO antibodies into reagent-friendly antibodies with rabbit constant regions, maintaining the human heavy and light chain variable domains. This allows use of secondary reagents to rabbit IgG to stain human as well as rodent tissues. Sections from mouse embryonic day 13 (E13) spinal cord or E16 telencephalon stained with a ROBO1 or a ROBO2 rabbit chimera antibody showed clear immunostaining patterns similar to those obtained with previously validated commercial antibodies (Supplementary Fig. S8). Both rabbit chimera IgG antibodies displayed reasonable off rates in SPR, with a k_d_ = 1.3 × 10^−3^ s^−1^ for the ROBO1 antibody and a k_d_ = 6.3 × 10^−^4 s^−1^ for the ROBO2 antibody (Fig. S8). Thus, the antibodies obtained from the minimal Fab library showed sufficient potency to be used as reagents.

### Deep sequencing analysis shows robustness and specificity of the ARM library

Deep sequencing revealed prominent clusters of related ARM sequences. Testing of the most populous member of each cluster revealed many potent antibodies. For instance, Cluster #3 for the IL23R antigen has 20 clones with sequence variation in the middle of the CDRH3 loop (Fig. 3A), indicating independent enrichment of related Fabs in the library and increasing confidence that these clones bind a specific epitope. Moreover, independent discovery campaigns for ROBO1 and ROBO2N, which are highly related with 94% sequence identity, showed enrichment of prominent clusters with essentially identical ARM sequences (Fig. 3B). Across the ten test antigens, CDRH3 diversity was reduced from 1,000 enriched CDRH3 clusters in the first FACS round (Fig. 3C) to only ∼100 clusters in the third FACS round. The number of sequences in each CDRH3 cluster increased to an average of 3 to 8 by FACS3 (Fig. 3D). Overall, the antibodies that were produced based on the most populous sequence cluster for each antigen after the final FACS round performed fairly well (17 out of 20 Fabs showed robust antigen binding (Supplementary Table S4)), but the best performing antibodies were derived from clusters with lower abundance (Supplementary Fig. S6) suggesting that there was no clear correlation between the abundance of an Fab clone and the potency of the resulting antibody.

**Figure 3.**
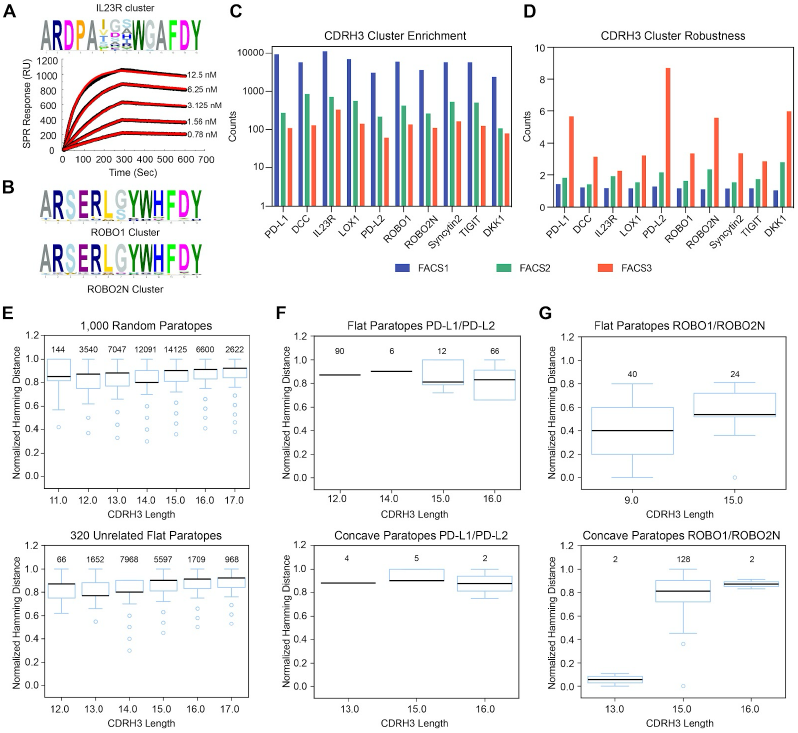
Antigen driven sequence enrichment shows robust and unique CDRH3 sequence patterns. **(A)** Sequence logo of the CDRH3 region of enrichment cluster 3 for antigen IL23R, showing high sequence consensus at the base of the CDRH3 loop and variability at the tip of the loop. The surface plasmon resonance sensogram is shown for the IgG1 antibody derived from the most dominant clone of this cluster **(B)** Comparison of one of the prominent CDRH3 clusters observed independently for the ROBO1 and ROBO2N antigen, indicating convergence of these enriched clusters. **(C)** CDRH3 cluster enrichment is shown over the three FACS selection rounds for each antigen. Sequences are clustered when they share more than 80% similarity according to the Blosum62 substitution matrix. Cluster counts are shown on a logarithmic scale. **(D)** CDRH3 cluster robustness is displayed as the average number of CDRH3 sequences that contribute to each cluster. **(E)** Normalized Hamming distances for each CDRH3 length for a random set of 1000 sequences from the unchallenged synthetic sequence dataset, and for all recombinantly generated antibodies with a flat paratope that are not targeting the same antigen. **(F)** Normalized Hamming distances for CDRH3s for antibodies generated for PDL1 and PDL2 with flat and concave paratopes. **(G)** Normalized Hamming distances for the CDRH3s for ROBO1 and ROBO2N generated antibodies with flat and concave paratopes. The number of pairwise sequences analyzed are listed for each CDRH3 length. Boxes indicate the interquartile range (IQR; Q1 to Q3), with the median shown as a horizontal line. Whiskers extend to the most extreme data points within 1.5×IQR from the quartiles; points beyond are plotted as outliers.

Serial enrichment for antigen binding is an imperfect process, as factors beyond affinity for antigen affect enrichment, including the efficiency of yeast display, heavy/light chain pairing, and differences in proliferation rates during yeast clone amplification^26^. We therefore examined if deep sequencing at each enrichment step provided additional information that could be exploited to obtain additional antibody paratopes. We first wanted to assure that the CDRH3 clusters obtained for a specific antigen were unique. Therefore, the sequence kinship between CDRH3 clusters selected for individual antigens was assessed by calculating their paired, normalized Hamming (NH) distances. CDRH3 sequences that have closely related amino acid sequences will have an NH distance approaching 0, whereas CDRH3 sequences that have a different amino acid in every position will have an NH distance of 1. NH distance analysis for a random set of CDRH3 sequences from the unchallenged library sequence data set showed that the median NH distance hovered around 0.85 for each of the designed CDRH3 lengths (Fig. 3E). When CDRH3 sequences from unrelated antigens were compared, most NH distances were as high as for the unchallenged library (Fig. 3E lower panel). PD-L1 and PD-L2 are two receptors with similar domain structures, which share 50 % sequence identity for the ectodomain construct used in antibody discovery (Fig. 2A). NH distance calculations revealed no overlap between the CDRH3 sequences obtained for PD-L1 and PD-L2 (Fig. 3F). This result aligned with the distinct epitope features for PD-L1 and PD-L2, which lack evolutionary conserved sequence patches on their surfaces

We also included two antigens (ROBO1 and ROBO2) that are closely related, with a sequence identity of 94 % for the overlapping epitope consisting of the N-terminal domain. The entire ectodomain of ROBO1 was screened, while for ROBO2, the antigen was truncated to the N-terminal immunoglobulin domain alone (Fig. 2A), involved in binding to the ligand Slit^27^. It was expected that several antibodies obtained for the truncated ROBO2 antigen would be crossreactive to ROBO1. Indeed, the pairwise NH distances between sequences enriched for ROBO1 and the truncated form of ROBO2 (ROBO2N) revealed several related CDRH3 clusters (Fig. 3G). The flat-shaped paratopes for ROBO2 were dominated by a short 9mer CDRH3 motif that is shared with ROBO1. The most abundant sequence motif (GTWI) was identical for both targets, and the related antibody only showed moderate binding properties to ROBO2 (Supplementary Table S4). The flat paratope ARM repertoire manifested other CDRH3 motifs for ROBO1, and the dominant paratopes in the 15-mer cluster were quite distinct with superior binding kinetics compared to the short motif. The NH analysis showed that the minimal Fab library provides similar antibody paratopes for antigens with high sequence identity, but that the diversity is high enough to create antigen-specific paratopes that do not overlap with other unrelated antigens.

### Exploiting the diversity of early selections to discover additional antibodies using machine learning

We trained a logistic regression (LR) model on the ROBO2N dataset (Fig. 4A), in which ARM motifs from early sorting rounds were scored according to their enrichment counts in MACS and FACS1, and k-mers were assigned weights based on their relative abundance in FACS1 relative to MACS (Fig. 4A). To validate the ROBO2N LR model, we assessed its performance to rank ROBO1 antibodies that were experimentally verified in cell display (Fig. 4C). If the model performed well, it should be able to distinguish ROBO1 antibodies that share paratopes with the ROBO2N ARM data set. Indeed, the ROBO2N LR model identified 15 ROBO1 antibodies with high scores based on the ROBO2N training set (Fig. 4C). All 15 ROBO1 antibodies bound the ectodomain of ROBO1, as confirmed by cell display assays. Importantly, the three ROBO1 antibodies that failed to bind ROBO2N in cell display, as well as the other 9 ROBO1 antibodies, had low regression model scores for ROBO2N. In contrast, a logistic regression model trained on ROBO1 ARM data gave high scores for all these antibodies, while the ROBO2N model assigned low scores (Fig. 4C, Supplementary Fig. S9), indicating that these antibodies are truly represented in the ROBO1 but not the ROBO2N experimental data set.

**Figure 4.**
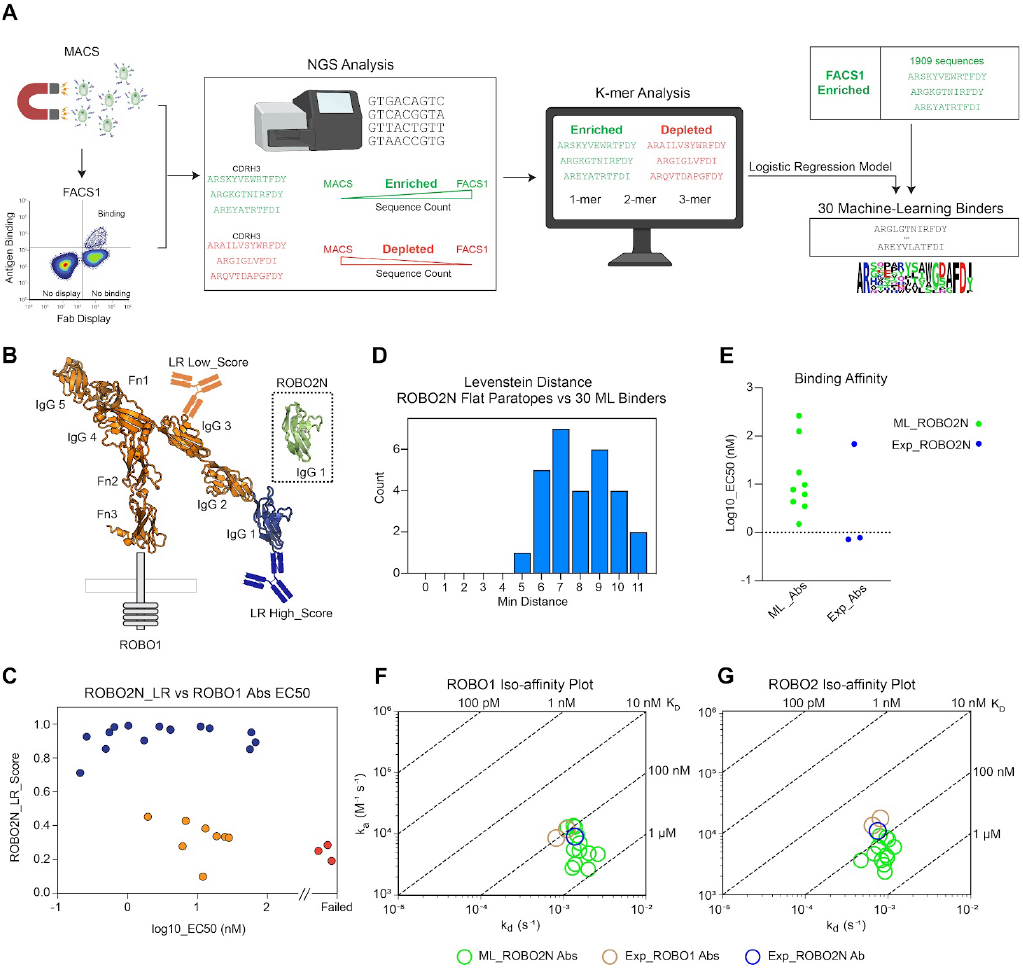
Logistic regression model of early sorting populations rescues ROBO2N binders lost during kinetic cell sorting. **(A)** Schematic of the logistic regression model used to identify promising Fab clones from the early MACS and FACS1 rounds, using k-mer frequences **(B)** The ROBO2N_LR model trained on ARMs binding to the N-terminal domain of ROBO2 was used to predict binding properties for the experimentally derived ROBO1 antibodies **(C)** Model benchmarking versus experimentally verified ROBO1 antibodies. Correlation between the ROBO2N_LR score (ranging from 0 to 1) and the cell display EC50 values experimentally obtained for the ROBO1 antibodies. Antibodies that failed to bind in cell display are colored red, antibodies that have similar scores for the ROBO2N_LR and ROBO1_LR models are colored blue and antibodies that have diverging scores between the ROBO2N_LR and ROBO1_LR models that are larger than 0.3 are colored yellow. **(D)** Histogram of closest Levenstein distances between the experimental and ML-derived ROBO2N antibodies. **(E)** Cell display antibody titration EC50 values for the ML derived ROBO2N antibodies (green) and the experimentally derived antibodies (in blue). **(F-G)** Iso-affinity plot of the ML derived ROBO2N antibodies and the ROBO1 and ROBO2 antibodies tested in IHC in human/rabbit chimera format using the ROBO1 **(F)** and ROBO2 **(G)** antigen.

Next, we scored all 1909 ARMs for ROBO2N from the first FACS round that had a significant sequence count (n>5). We selected 29 high-scoring ROBO2N ARMs that were depleted from later FACS selection rounds, and therefore overlooked during the experimental campaign (Supplementary Table S5). To ensure novelty, the selected sequences had a Levenshtein distance of at least five from any previously ordered ROBO1 or ROBO2N antibodies (Fig. 4D). We used the Levenshtein distance instead of Hamming distance, to rule out similar sequences across different lengths. The 29 ML-derived ARMs were converted to antibodies with rabbit constant regions, which is the isotype used for reagent format and tested for binding properties using SPR and cell display. Nine out of the top ten ML-derived antibodies exhibited strong binding kinetics comparable to the antibodies for ROBO1 and ROBO2 that performed well in our IHC experiment (Fig. 4E and 4F), and with superior off-rates compared to the dominant identical ROBO1 ROBO2N clones with a shortened CDRH3 sequence (Supplementary Table S4 and S5). Titration of the antibodies onto cells displaying ROBO2 confirmed binding for eleven antibodies (Fig. 4E). Crossreactivity tests by SPR demonstrated that all 11 ML-derived ROBO2N antibodies also bound ROBO1, which was to be expected due to the high sequence identity between these antigens. In contrast, no off-target crossreactivity was observed for negative control antigens, including PD-L1, PD-L2, DCC and LOX1. We applied the same approach to our PD-L2 ARM flat paratope data set, which had only given two good binders experimentally. The logistic regression model scored 33 ARM motifs with sufficient Levenshtein distance from the early FACS round, and found that 17 antibodies showed comparable potency in cell display, although the SPR properties were somewhat diminished (Supplementary Table S6 and Supplementary Fig. S10). Together, the expansion of antibody candidates for ROBO2N and PDL2 through this machine learning approach highlights the underlying richness of the deep sequenced ARM data sets.

## Discussion

Advances in machine learning enabled significant progress in predicting protein complexes at scale from sequences alone but required deep sequence alignments to detect coevolution^28,29^. More recent work has demonstrated some success in predicting interactions when learnt from known 3D structures including antibody protein interactions^30,31^ and even design^31,32^. However, this in *silico* prediction of antigen specific antibodies has so far been limited to redesigning antibody-antigen interfaces for which rich 3D structural information is available. Synthetic libraries have the potential to generate robust antibody/antigen data sets that could be used for machine learning applications^33^, but these data sets have so far been limited in scope and contain a variety of binders that rely on contributions of all CDR regions. The library presented here constrains sequence space, focusing molecular diversity primarily within the CDR3 region of the heavy chain. The readout of the antigen binding features was encoded in a compact antigen recognition module (ARM) well-suited for machine learning applications. We tested this library in a parallel discovery campaign on a diverse range of antigens. Most antibodies obtained from this discovery campaign contained paratopes represented by a cluster of related amino acid sequences, from which a single sequence was selected, underscoring the robustness of the Fab clone selection process and the uniqueness of each paratope for its respective antigen.

While the cell sorting procedure resulted in many antibody candidates, it became clear that the loss of clonal diversity was not compensated by necessarily obtaining all the best binders, a limitation of display technologies that has been observed previously^26,34^. We therefore exploited the simplicity of the ARM design to reevaluate the larger pool of clones available from the deep sequencing of the early selections for ROBO2 and PDL2. Using a logistic regression model, we were able to significantly expand the number of antibody candidates for these antigens, showcasing the promise of a hybrid in silico/experimental antibody discovery approach.

The extensive NGS data accompanying these discovery campaigns provide a solid foundation for developing machine learning models for *in silico* epitope binning, affinity maturation and ultimately, zero shot antibody discovery. To facilitate future advancements in machine learning-driven antibody discovery, we have made the complete sequencing data set available, along with the antibody characterization for 486 antibodies (See Supplementary data and materials). This data set includes sequences for each antigen, enrichment data across selection rounds, and information on non-binding clones, offering valuable entry points for future antibody discovery efforts. The data set contains more than 68,000 unique ARM sequences with at least two ARM counts associated with their target antigen. It should be noted that this dataset may require processing to remove noise resulting from PCR or other sequence errors, as well as crossover of low abundant clones between antigens as a result of crosscontamination during the lysis of the yeast cell pools.

The primary goal of this library is to enable high-throughput discovery of reagent antibodies targeting cell surface receptor families for research applications that can also serve as leads for therapeutics. Previous iterations of this library have been successfully used to develop antibodies against RGD-binding integrins^35,36^, where FACS selection identified antibodies with specificity for certain alpha/beta integrin pairs. The antibodies generated with this library design have been validated using rigorous methods typically applied to therapeutic antibody development and demonstrated suitability for various applications, in particular flow cytometry and immunohistochemistry. Future challenges include an expansion of the library to cover other heavy/light chain pairs, an increase in throughput and ML efforts to systematically relate the biophysical properties of these antibodies to their performance across a broader range of antibody reagent applications. Our aim is to establish a large-scale effort to provide reliable antibody reagents to the broader research community^37^ while advancing an Open Science approach to antibody discovery that integrates machine learning with high-throughput experimental validation.

## Supplementary Data

NGS sequencing files and a library key file are provided in csv format along with this manuscript. The NGS data contain all unique CDR3 sequences obtained after FACS1, FACS2 and FACS3, and n>0 counts per sequence for the ROBO1, ROBO2N and PDL2 MACS1 data and is deposited in Zenodo: https://zenodo.org/records/15367597. The 486 experimentally tested antibodies are provided along with biophysical characterization information.

The code to weigh and classify k-mers using logistic regression has been deposited at the Github repository.

## Supplementary Materials and Methods

This manuscript is accompanied by nine supplementary tables and ten supplementary figures.

## Materials and Methods

### Generation of the Fab library

#### Construction of VH1-69 Fragments and Pairing with Light Chains

The VH1-69 heavy chain was constructed entirely using germline sequences. For the CDRH3 region, the first two residues conformed to the consensus A105R106, while the final residues followed the consensus J region sequence F115D116Y117, with exceptions for lengths of 11 (F/LDY) and 15, 16, and 17 (FDY/I/P/V). Position-specific amino acid frequencies for the intervening CDRH3 positions were introduced through combinatorial variant oligo synthesis, mimicking the naïve CDRH3 repertoire (OAS database) while avoiding known sequence liabilities. Cysteine and methionine residues were excluded to prevent oxidation liabilities. CDRH3 lengths ranged from 11 to 17 amino acids, reflecting the most productive and dominant lengths in the naïve repertoire.

Four VH1-69 fragments were generated, each incorporating synonymous substitutions in the J and CH1 regions (light chain barcode). These fragments were paired with four distinct light chains. Prior to library construction, four yeast strains were created by transforming each strain with a single light chain expression plasmid. The light chain plasmids contained VK1-39, VK3-15, VK3-20, and VK4-1, each featuring fully germline CDRL1 and CDRL2 loops. The CDRL3 sequence reflected a common VJ rearrangement for the respective V gene, with a resulting CDRL3 length of 9. These clones were expanded for large-scale heavy chain transformations, with each VH-VL pair forming a unique sub-library. VK4-1, featuring a longer CDRL1 than the other light chains, contributed to the creation of a concave paratope known for its affinity for linear epitopes or peptide antigens^18,19,25^.

#### Library Construction and Transformation

To construct the Fab library, the four yeast strains containing different light chains were transformed with a heavy chain base vector linearized between the signal sequence and constant region, along with VH1-69 fragments containing CDRH3 diversity and the corresponding light chain barcode. Both heavy and light chains were expressed from a CEN vector with TRP1 and URA3 selection markers, respectively. Expression was controlled by the GAL1 promoter, and both chains featured an N-terminal *S. cerevisiae* alpha-factor signal sequence for secretion. The heavy chain was fused to Aga2 at the C-terminal side and further tagged with a V5 epitope for detection. Light chain expression was detected using a c-myc tag placed C-terminally to the VL. Over 4 × 10^9^ transformants were generated across the four VH/VL pairs. The library was archived by suspending approximately 1 × 10^10^ cells (10X coverage) for each VH-VL pair in 1.8 mL freeze media (URA TRP dropout media with 15% glycerol) in 2 mL cryogenic vials and stored at -80°C.

### Strain for yeast display

The genomic copy of AGA2 was deleted from *JAR200* (MATa, GAL1-AGA1::KanM×4ura3D45, ura3–52 trp1 leu2D1 his3D200 pep4::HIS3 prb1D1.6R can1) using one step PCR mediated deletion to replace the AGA2 ORF with the hphNT1 gene that provides resistance to hygromycin.

### Yeast Electroporation

The yeast electroporation procedure followed the established protocol outlined by Benatuil et al., 2010^39^, with slight modifications. The library was created in the *S. cerevisiae* strain *JAR200 aga2hphNT1*. A 200 ml yeast culture at an OD of 1.6/ml was used for each electroporation. Prior to electroporation, cuvettes were pre-chilled at -20°C. After electroporation, cells were recovered in 10 ml of a 1:1 mix of 1 M sorbitol and YPD media. The cells were then incubated at 30°C for 1 hour with shaking at 120 rpm before dilution plating onto SD-URA-TRP plates to calculate library size. The remaining cells from the electroporation reaction were cultured in 200 ml of SD-URA-TRP broth in a baffled flask with agitation at 225 rpm and 30°C, forming the transformed library.

### Antibody discovery using MACS and FACS

A yeast display library comprising over 3 × 10^9^ transformants for flat paratopes VH1-69/VK3-15, VH1-69/VK3-20 and VH1-69/VK1-39 and 1 × 10^9^ transformants for concave paratopes VH1-69/VK4-1 was used each for separate “flat paratope” and “concave paratope” campaigns. For MACS, the unchallenged library was grown in -URA -TRP dropout media for 2 days and Fab expression was induced for a duration of 24 hours in SGCAA media. On the day of the MACS experiment, 100nM biotinylated antigen was incubated with 500 µl of streptavidin microbeads (130-048-101) in a total of 1 ml of M buffer (20 mM HEPES, 150 mM NaCl, 2 mM CaCl2, 2 mM MgCl2, 20 mM maltose and 0.1% BSA) for one hour at 4°C. Subsequently, this mixture was incubated with the library pool in 25 ml of M buffer for 30 minutes at room temperature and processed through the Miltenyi autoMACS pro system (130-092-545) using the posslD2 protocol. Cells obtained from the MACS elution were grown for two days in -URA -TRP dropout media, followed by induction in SGCAA for 24 hours, and then subjected to FACS.

Before the first round of FACS a negative selection against streptavidin microbeads was performed. 100 ul of streptavidin beads were incubated with 1× 10^8^ yeast cells in 2 ml of M buffer for 30 minutes at room temperature followed by processing on autoMACS using posslD2 protocol. An additional depletion against 100nM biotinylated Fc was conducted in case of Fc-fused antigen. The flow-through, containing the majority of the 1 × 10^8^ yeast cells, was incubated with 100nM biotinylated antigen in M buffer for one hour at 4°C. After washing with M buffer, the cells were stained with a 1:100 dilution of Alexa Fluor 405-conjugated anti-V5 antibody (homemade) to detect heavy chain expression, a 1:100 dilution of FITC-conjugated anti-myc antibody (Immunology Consultants laboratory-CMYC-45F) to detect light chain expression, and a 1:200 dilution of APC-conjugated streptavidin (TONBO Bioscience-204317U100) for 10 minutes at 4°C. Following a wash with M buffer, the cells were resuspended in M buffer for FACS using a SONY SH800 cell sorter. Gates were drawn to identify paired Fab expression and antigen-binding clones, which were then sorted. In the second round of FACS, cells were incubated with 20nM biotinylated antigen for one hour at 4°C followed by a wash and subsequent incubation with 200nM unbiotinylated antigen. For the last round of sorting, antigen concentration was dropped to 4nM. A final analytical stain was conducted to confirm antigen binding before subjecting the sorted population to NGS. Flow cytometry gating figures were generated using FlowJo.

### Novaseq library preparation

Frozen vials containing 1 × 10^10^ cells of each VH1-69/VK3-15 and VH1-69/VK3-20 libraries were inoculated in 1L of SD -TRP -URA growth media and grown overnight at 30 with shaking at 200 rpm. 1 × 10^10^ cells from the overnight grown saturated yeast cultures were then aliquoted across 96 well deep 2mL plate. Cells were spun down at 2400xg for 3 min, resuspended in buffer containing Zymogen’s Zymolyase lytic enzyme and incubated for one hour at 37, shaking at 1000 rpm. Following zymolyase treatment plasmids were extracted using Qiagen PLASMID Plus 96 kits per manufacturers protocol. For deep sequencing an initial round of PCR was performed on eluted plasmid from ∼ 2 × 10^9^ cells using forward primer that binds in FR2 and reverse primer that binds in CH1 (Table S7) to amplify the VH1-69 gene from the heavy chain plasmid. The entire PCR product was then pooled together and purified using 0.8x AMPure SPRI bead cleanup (Beckmann Coulter). A second round of PCR was then performed across a 96 well plate for Illumina Nextera Adapter annealing and barcode addition. PCR products were pooled together and purified through two rounds of 0.8X AMPure SPRI bead cleanup. Concentration of the final pooled library was measured using Qubit Fluorometric Quantification and the final library was spiked with a 10% PhiX as a control. Sequencing was performed on Illumina’s Novaseq X platform.

### NGS of selection outputs

All selection outputs (selection rounds MACS, FACS1, FACS2, FACS3) were stored frozen in 15% glycerol at -80°C. Frozen aliquots were inoculated in SD -trp -ura growth media (seeded at about 1 OD/mL) and grown over night in 30°C incubator shaking at 200 rpm. Next day, cells were spun down at 2400xg for 3 min and resuspended in buffer containing Zymogen’s Zymolyase lytic enzyme and set to incubate for one hour at 37°C, shaking at 1000 rpm. Following zymolyase treatment, plasmids were extracted using Qiagen PLASMID Plus 96 kits per manufacturers protocol. For deep sequencing an initial round of PCR was performed on eluted plasmids using a forward primer that binds in FR2 and reverse primer that binds in CH1 (Table S7) to amplify the VH1-69 gene from the heavy chain plasmid. A second round of PCR was then performed across a 96 well plate for Illumina Nextera Adapter annealing and barcode addition. PCR products were pooled together and purified through two rounds of 0.8X AMPure SPRI bead cleanup (Beckman Coulter). The concentration of the final pooled library is measured using Qubit Fluorometric Quantification. Library samples were diluted to 4nM and denatured following Illumina Miseq System Denature and Dilute Libraries Guide. Final library concentration of 12pM with a 10% Phix spike in control was run on an Illumina Miseq platform.

### MiSeq and NovaSeq data analysis and antibody clustering

Paired sequencing reads were extracted from the Illumina NovaSeq or MiSeq systems and merged using BBMerge^40^ to create consensus nucleotide sequences across the amplicons. The assembled sequences were subsequently trimmed to the PCR primers used in sample preparation. Light chain information was determined using the barcode incorporated in the sequences. The DNA sequences were further translated for CDR3 extraction. Only sequences having exact matches to the VH1-69 backbone were retained for further analysis.

For antibody discovery, results from each cell sorting selection round were grouped based on the antigen used and unique CDR3 sequences and ranked by sequence abundance found in the final selection round. Similar sequences were clustered using the following centroid-based method: The most abundant CDR3 sequence from each selection round was automatically assigned as cluster #1. Each subsequent CDR3 sequence was compared to sequences of higher abundance. For a CDR3 sequence of identical length as the reference, a sum of the pairwise comparison scores using BLOSUM62 table was calculated. If the sum was greater than 80% of the maximum (pairwise comparison of the reference sequence to itself), the CDR3 was then assigned the same cluster number as the reference. If the CDR3 showed no similarity to any of the previous clusters, it was assigned a new cluster number. This process was repeated for all CDR3 sequences that had at least one read count. After clustering, we selected the VH sequences with the most abundant CDR3 from each cluster and produced these as antibodies for characterization.

### Antigen production

Sequences encoding predicted mature secreted glycoproteins and integral membrane glycoprotein ectodomains (Table S8) were codon optimized, synthesized and cloned in the pcDNA3.4-TOPO mammalian expression vector (Thermofischer) by Thermo/GeneART, SynBio or Twist Biosciences. All constructs were fused with an N-terminal secretion signal peptide (Table S8) and C-terminal Avi tags for biotinylation and 10X Histidine tags for affinity purification^41^. In Lox-1, a single-pass type II transmembrane protein, the tags were fused to the N-terminus after the signal peptide. Additionally,the IL-23 construct was fused with a C-terminal human IgG1 Fc fragment in addition to the tags described above, to improve yield and stability.

For the expression of antigens in mammalian cells, Gibco™ Expi293F™ cells (Thermo Fisher Scientific) were diluted to a concentration of 3×10^6^ cells/mL 24 hours prior to transfection, allowing them to double overnight at 37°C. At the time of transfection, the cells were again diluted to 3×10^6^ cells/mL. The DNA was prepared by diluting it to 0.8 µg/mL in 1/10th the transfection volume using Gibco™ Opti-MEM™ Reduced Serum Medium (Thermo Fisher Scientific), and then complexed with 0.8 µL/mL of Fecto-PRO® DNA transfection reagent (Polyplus) for 10 minutes. This mixture was added to the diluted Gibco™ Expi293F™ cells in the transfection flasks. The transfected cells were incubated in a shaker incubator at 37°C for 24 hours. Protein expression was enhanced by adding Valproic acid sodium salt (VPA, Sigma-Aldrich) and D-(+) Glucose (Sigma-Aldrich) to final concentrations of 3 mM and 0.45% (v/v), respectively. After feeding, the flasks were returned to the incubator and grown for an additional four days at 32°C.

Secreted antigens were harvested by centrifuging the culture at 5000 x g in 50 mL conical tubes. For antigen purification, 1 mL of Nickel beads slurry (cOmplete His-Tag Purification Resin, Roche) pre-equilibrated in 20 mM HEPES pH 8, 250 mM NaCl, 1 mM CaCl_2_, and 1 mM MgCl_2_ (Binding Buffer) were added to the 50 mL of harvested antigens and incubated for 1 hour at 4°C on an SRT roller (Cole-Palmer). The bound antigens were then purified by gravity flow using Poly-Prep® chromatography columns (Bio-Rad). The antigen and nickel bead mixture were loaded into the chromatography column, allowing the beads to settle. Residual beads were collected by washing the conical tubes with 10 mL of Binding Buffer. The packed nickel beads were then washed twice with 10 mL of 20 mM Imidazole in Binding Buffer. Antigens were eluted following a 5-minute incubation with 2.6 mL of Binding Buffer containing 250 mM Imidazole. Eluates were further purified by size exclusion chromatography on a GE HiLoad 16/600 Superdex 200 pg column using an AKTA Pure 25 L (GE Healthcare). Antigens were eluted in the buffer 20 mM HEPES pH 7.4, 150 mM NaCl, 1 mM CaCl2, and 1 mM MgCl2. All buffers were supplemented with 1 mM TCEP, except when used for purification of the Fc tagged IL-23.

The purity and integrity of the purified antigens (ensuring no cleavage products) were verified by SDS-PAGE. Additionally, monodispersity and stability were assessed by Dynamic Light Scattering (DLS) following a single freeze-thaw cycle. Antigens that did not aggregate were biotinylated using either *E. coli* Biotin Ligase (BirA) or EZ-Link™ Sulfo-NHS-LC-Biotin (Thermo Fisher Scientific). For enzymatic biotinylation, 10 mM ATP and 10 mM magnesium acetate were added to the antigen, followed by biotin at a concentration 10 times the total molar amount of antigen, and 0.3 µM BirA per 100 µM antigen. The reaction mixture was incubated at 4°C for 18 hours. For chemical biotinylation, the antigen was incubated with EZ-Link™ Sulfo-NHS-LC-Biotin at a concentration 10 times the total molar amount of antigen, also overnight at 4°C. The following day, both reactions were processed to remove any unreacted biotin, ATP, and magnesium acetate using Zeba™ Dye and Biotin Removal Spin Columns (Thermo Fisher Scientific), followed by filtration through a Vivaspin 0.2 µm spin filter.

Biotinylation levels were evaluated based on the migration patterns of biotinylated antigens in the presence of Streptavidin. For the assay, 1 µg of biotinylated antigen was incubated with 1 µg of Streptavidin (New England Biolabs) for 20 minutes on ice. Subsequently, 2-5 µL of 4x Laemmli Sample Buffer (lacking reducing agent) was added to the antigen-Streptavidin mixture. The samples were run on an SDS-PAGE gel, with a control sample of the antigen (without Streptavidin) loaded alongside. A slower migration of the biotinylated antigen in the presence of Streptavidin indicated successful biotinylation. To quantify the biotinylation efficiency, the intensity of the protein band corresponding to the antigen’s molecular weight was compared between the lanes with and without Streptavidin. Biotinylation was considered successful if the intensity of the band in the Streptavidin-containing lane was 30% or less of that in the control lane without Streptavidin. Biotinylated antigens were tested again for monodispersity and stability using DLS as described above. Stable antigens were mixed 0.005% (w/v) Tween-80 (cryo-protectant), diluted to 4 µM, aliquoted to 100 µL and flash frozen in liquid nitrogen and stored at -80°C.

### Antibody production

#### Antibody plasmid construction

The entire variable region of VH1-69 (IMGT1-128) including the CDRH3 were synthesized and inserted into a pcDNA3.4 expression plasmid that already contains the constant domains of human IgG1 (Human IgG1_CH1CH2CH3). These were paired with fixed light chains (Human VK1-39, Human VK3-15, Human VK3-20, and Human VK4-1), based on sequencing results. For conversion to the human/rabbit chimera, the variable domains of the heavy chain (IMGT1-128) were fused with the constant region of rabbit IgG (Rabbit_IgG_CH1CH2CH3). The variable domains of the light chains were fused with the constant region of the rabbit kappa light chain for matching Human/Rabbit_VK1-39, Human/Rabbit_VK3-15, Human/Rabbit_VK3-20, and Human/Rabbit_VK4-1. Each individual antibody candidate is barcoded and annotated in a 96-well plate format for data tracking. Heavy chain plasmids of antibody candidates are synthesized by Twist Biosciences in 96-well plate format. The heavy chain DNA is resuspended in molecular biology grade water (IBI Scientific) to a final concentration of 100 ng/µL using a Biomek i7 automated workstation (Beckman Coulter). The light chain DNA is prepared with a Giga prep kit (Machery-Nagel) following the manufacturer’s protocol. To prepare for transfection, heavy chain DNA and the corresponding light chain DNA are mixed by the Biomek i7 to a final concentration of 25 ng/uL and 62.5 ng/uL, respectively, in another 96-well titer plate (Greiner Bio-One).

#### High-throughput antibody production

On the day of transfection, Expi293 cells (ThermoFisher) are seeded at 3e6 cells/mL in Expi293 expression medium (ThermoFisher) with viability above 90%. In brief, 23 µL of the premixed DNA, 2 µL of FectoPRO transfection reagent (Polyplus), and 180 µL of serum-free Opti-MEM media (Gibco) are mixed and incubated on a Biomek i7 at room temperature for 10 minutes in a 96-well titer plate to allow complex formation. The transfection mixture is split into two 96-well deep-well plates after incubation and 1 mL of seeded Expi293 cells are added to each well. This results in a total of 2 mL expression culture per selected antibody candidate. The transfection plates are sealed with breathable silicone adhesive seals (Analytical Sale and Services) and incubated at 37 ºC, 1000 rpm, 8% CO_2_, at 80% humidity (INFORS HC, Multitron Pro). The transfection plates are supplied after 24 hours with valproic acid sodium salt (VPA, Millipore Sigma) and D-(+) glucose (Millipore Sigma) to a final concentration of 3 mM and 0.45% respectively. The transfection plates are then sealed and returned to the incubators and allowed to grow for another 6 days under the same incubation conditions.

The fully grown cultures are harvested by centrifuging at 3,200 g for 30 minutes at room temperature. Supernatants are then collected from the transfected plates to 96-well deep well plates (Greiner Bio-One) using Biomek i7. To purify the antibodies, 25 µL of Protein A magnetic beads (Lytic solutions) are added to each well of the supernatant and plates are incubated at 37 °C, 1000 rpm (INFORS HC, Multitron Pro) for 2 hours to allow capturing. Antibody plate purifications are performed on a Biomek i7. To summarize, supernatants are transferred to a waste reservoir (Agilent) with a 96-well magnet (Alpaqua) separator at the bottom. Then the beads are washed sequentially with 500 µL/well of the bind buffer (25 mM HEPES, 150 mM NaCl, pH 8.0), and 500 µL/well of the wash buffer (25 mM HEPES, 1 M NaCl, pH 8.0). In each step of the wash, magnetic beads are fully suspended for 10 minutes via pipetting and the washing liquid is discarded to the waste reservoir with magnet separation. Finally, beads are subjected to 90 µL/well of the elution buffer (100 mM Glycine, 150 mM NaCl, pH 3.0) and a 5-minute, 1,000 rpm incubation (Bioshake) at room temperature. The elution and beads are then transferred to a 96-well filter plate (Thompson) for final clearance. The filter plate is stacked on top of a 96-well titer plate that contains 10 µL/well of the neutralization buffer (1 M HEPES, pH 8.0), and the final products are collected by centrifugation at 1,000 g for 2 minutes. The final products are mixed and then loaded into high Lunatic plates with the Biomek i7 for concentration assessment. The antibody concentrations are quantified using the common A280 application on a Lunatic Spectrophotometer (Unchained Labs) with an extinction coefficient of 13.7. Antibodies with yields less than 50 mg/L are marked as “Fail” in purification.

#### High-throughput antibody quality analysis

Purified antibodies are all normalized to a concentration of 0.5 mg/mL in 1xPBS buffer pH 7.4, and then regrouped in 384-well plates (USA Scientific). Up to 23 antibodies per antigen are rearranged in the same row, and up to 16 discovery campaigns are combined in a 384-well plate. The 24th column of each 384-well plate contains 4 replicates of the following monoclonal antibody controls: 4E10, cixutumumab, basiliximab, and adalimumab. The polyspecific reactivity (PSR) of the antibodies is evaluated by a single-concentration enzyme linked immunosorbent assay (ELISA). The PSR-ELISA experiment is fully automated with a Biomek NXP automated workstation (Beckman Coulter)-centered integrated system. Generally, 4 maxi-sorp 384-well plates are coated with four substrates, namely 0.1 µg/well of bovine genomic DNA (Amsbio), 0.02 µg/well of insulin (Millipore Sigma), 0.2 µg/well of SF9 surface membrane preps (SMP), and 0.1 µg/well of avidin (Millipore Sigma), respectively, and blocked by 3% BSA afterward. Antibodies are diluted to a final concentration of 10 µg/mL and screened for binding to these substrates. Alkaline phosphatase goat antihuman IgG (Jackson ImmunoResearch Laboratories) is used for detection at a dilution of 1:5,000 to a final antibody concentration of 0.12 µg/mL. In between steps, plates are washed 5 times with 1xPBST buffer pH 7.4 using a microplate washer (BioTek). Color development is performed with a 30-minute p-Nitrophenyl phosphate substrate (Millipore Sigma) incubation, followed by immediate quenching with 5 M sodium hydroxide (Millipore Sigma). Finally, raw data are collected using a SpectraMax i3x at 405 nm. All readings are normalized with respect to basiliximab. Antibodies are considered as “Fail” in the PSR assay when showing higher affinity than basiliximab against more than 2 of the substrates tested.

The analytical size-exclusion chromatography (SEC) assessment is performed with an Agilent 1260 Infinity II isocratic pump system. 10 µL of each antibody at 0.5 mg/mL is transferred to a 384-well PCR plate (Bio-rad) using Biomek NXP automated workstation and sealed with a pre-slit adhesive seal (Analytical Sale and Services) to prevent evaporation. Antibody samples are analyzed with a TSKgel SuperSW mAb HTP column (Tosoh, 4um 1.6mmx15cm) at a wavelength of 280 nM with a flow rate of 0.5 mL/min in 1xPBS buffer pH 7.4. Routine column cleaning cycles are performed after every 96 samples following manufacturer’s recommendations. Antibodies with multiple detectable peaks (dominant peak area <90%) or with out-of-range retention times are characterized as “Fail” in SEC.

#### Medium-scale antibody production

Similar to small scale antibody production, Expi293 cells are seeded at 3e6 cells/mL prior to transfection. Per antibody transfection, a mixture is prepared of 6.86 µg of the heavy chain plasmid, 17.14 µg of the light chain plasmid, 24 µL of FectoPRO transfection reagent, and 2 mL of serum-free Opti-MEM media in a 50 mL culture tube (Midland Scientific). Each mixture is incubated at room temperature for 10 minutes and then added to 28 mL of media containing 3e6 Expi293 cells. Expression cultures are grown at 37 ºC, 1000 rpm, 8% CO_2_, and 80% humidity (INFORS HC, Multitron Pro) for 24 hours, and then fed with 3 mM of VPA and 0.45% D-(+) glucose.

Fully grown cultures are harvested 6 days after feeding. Each centrifugation-clarified supernatant is incubated with 0.5 mL of Protein A agarose resin (Lytic solutions) at room temperature for 2 hours to ensure sufficient binding. After incubation, the resin slurry is transferred to an empty column and to let the supernatant drain through. The resin is washed with 10 column volume (CV) of bind buffer, then 10 CV of wash buffer to remove nonspecifically bound contaminants. Following the wash, antibodies are eluted with 5 CV of elution buffer after 5-minutes of incubation and immediately buffer exchanged into 1xPBS (pH 7.4) using PD-10 columns (Cytiva). The concentration of the purified antibodies is measured by Nanodrop (ThermoFisher) using an extinction coefficient of 13.7, and then stored at 4 ºC for future usage.

### High throughput surface plasmon resonance

The SPR binding kinetics was measured using the Carterra LSA platform with HC30M sensor chips (Carterra, cat. #4279) at 25°C. Each chip was initially covalently functionalized via amine coupling with either goat anti-human IgG Fc antibody (Jackson ImmunoResearch Laboratories, cat. #109-006-098) or goat anti-rabbit IgG Fc antibody (Jackson ImmunoResearch Laboratories,cat. #111-005-046), depending on the capture antibody format. The sensor surface was activated by injecting a mixture of 0.033 M N-Hydroxysuccinimide (NHS, Millipore Sigma, cat. #130672) and 0.133 M N-(3-dimethylaminopropyl)-N’-ethylcarbodiimide hydrochloride (EDC, Millipore Sigma, cat. #E6383) in 0.1 M MES (Carterra, cat. #3625) buffer at pH 5.5 for 7 minutes. Following activation, the goat anti-human or anti-rabit IgG Fc was immobilized by injecting a 100 µg/mL solution in 10 mM sodium acetate, pH 4.5, for 10 minutes. Unreacted esters were quenched with 1 M ethanolamine, pH 8.5 (Carterra, cat. #3626) for an additional 10 minutes, resulting in approximately 5000 response units (RU) of immobilization.

Kinetic binding assays were conducted using an antigen buffer consisting of 20 mM HEPES pH 7.4, 150 mM NaCl, 1 mM CaCl_2_, 1 mM MgCl_2_, and 0.005% Tween 80. Tested antibodies were captured by pre-immobilized goat anti-human or anti-rabbit IgG Fc using a 96-channel print-head, and realtime kinetic measurements were performed by titrating the antigen from low to high concentrations. In specific cases, such as antigens fused to Fc domain (eg., hIL23R), we directly immobilized the tested antibodies onto the sensor chip to prevent binding from the Fc part of antigen, and performed kinetic measurements while regenerating in between antigen titration. The highest concentration of antigen started at 400 nM, followed by two-fold dilutions across eight subsequent concentrations to a minimum concentration of 3.125 nM. For each concentration, data collection times were set to 60 s for baseline, 300 s for association, and 300 s for dissociation. At the end of each antigen’s kinetic titration, the chip surface was regenerated using 0.85% phosphoric acid with three 30-second pulses.

The raw kinetic titration data were pre-processed in the Carterra Kinetics software, which included reference subtraction using inter-spots with no immobilized antibodies, buffer subtraction with the leading buffer cycle, and data smoothing. The kinetic data were then analyzed using a 1:1 Langmuir binding model with global fitting for both association (k_a_) and dissociation (k_d_) rates utilizing the Carterra kinetic software. The equilibrium dissociation constant (K_D_) was calculated as the ratio of the dissociation to association rate constants (K_D_ = k_d_/k_a_).

### Cell display assay

Expression plasmids (Synbio Technologies, GenScript Biotech, and TWIST Bioscience, see Table S9) were synthesized to display the proteins on the cell surface. For IL23R, ROBO1, ROBO2, PDL1, PDL2, TIGIT, DKK1, and DCC, the extracellular domains of both human and mouse orthologs were fused with a V5 tag amino acid sequence and the transmembrane domain from human PDGFR (AVGQDTQEVIVVPHSLPFKV<u>VVI-SAILALVVLTIISLIILI</u>MLWQKKPR). For Syncytin2, the full-length protein (16-538 A.A.) was used for making the expression construct. This is because the native receptor might form a trimeric complex through the cytoplasmic domains. For LOX1, which is a transmembrane type 2 protein, the extracellular domain is at the carboxyl end. The expression construct was made by fusing a human OX40L leader sequence (MERVQPLEEN-VGNAARPRFERNK<u>LLLVASVIQGLGLLLCFTYICLHFSAL</u>) including the signal peptide and transmembrane domain with sequences encoding the V5 tag and either the human or mouse LOX1 extracellular domain.

The expression plasmids were transfected transiently into ExpiCHO-S cells (Thermo Fisher Scientific) using ExpiFectamine CHO (Thermo Fisher Scientific) per manufacturer’s instructions. Before antibody titration, the expression levels were checked by applying an anti-V5 antibody (Thermo Fisher Scientific), to detect cell surface receptor expression, which was then detected by Alexa647-conjugated Rabbit anti-mouse secondary antibody (Jackson ImmunoResearch). The cell staining was assayed in an Intellicyt iQue Plus flow cytometer (Sartorius). The optimal transfection levels were determined by titrating the plasmid DNA used in transfection. In this way, we calibrated the detection of antibodies binding to the cell surface receptors without saturating the iQue system. Since the Syncytin2 constructs do not have a V5 tag, the maximum amount of DNA was used. Syncytin2 signals obtained from the actual antibody staining were well below the saturating levels of the iQue instrument.

For antibody titration, both human and mouse constructs were transfected using the optimal amount of DNA. For human genes, the cells are co-transfected with a GFP expression construct. For mouse genes, the cells are cotransfected with a BFP expression construct. Two days after transfection, cells were collected and blocked in 1% FBS in DPBS on ice for 30 min. After blocking, the cells were filtered through 40 um cell strainers and counted. Cells expressing human and mouse orthologs were mixed 1:1 and loaded in the 384-well plates using a Lynx LM900 liquid handling robot with a VVP96 head. Approximately 1 × 10^5^ cells were loaded into each well. All antibodies were serial diluted in DPBS from 660 nM (100 ug/ml) to 4.4 pM in 1:2.5 ratio using the Lynx LM900 robot in 384 well plates. Human IgG1 isotype control antibody (Bio X Cell) was also included in the assay at the concentration of 100 nM (15 ug/ml).

Antibodies were applied to cells using a BioMek NXP liquid handling robot with a multi-channel 384 head. All the subsequence washes and secondary antibodies applications were carried out in this system. Cells were incubated with primary antibodies for 30 min on ice and then washed twice in DPBS. Alexa647-conjugated donkey anti-human F(ab’)_2_ fragments (Jackson ImmunoResearch) diluted 1:200 in DPBS were then added as secondary antibodies. After incubation on ice for 30 min, the cells were washed twice in DPBS. The binding parameters of the antibodies to the cells were assessed according to the mean fluorescence intensities of Alexa647. The EC_50_ of the binding to human and mouse orthologs were determined by curve fitting using the following equation:

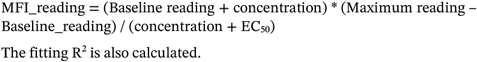

The fitting R^2^ is also calculated.

### LOX1 and TIGIT antibody benchmarking in cell display

The comparisons with commercial LOX1 and TIGIT antibodies carried out using the same cell display method described above with the following exception. The secondary antibodies used for the commercial antibodies were an Alexa647-conjugated Rabbit anti-mouse secondary antibody (Jackson ImmunoResearch). The same 1:200 dilution was used. The commercial antibody for hTIGIT is a mouse Anti-human TIGIT (clone TgMab-2, Becton Dickinson). The commercial antibody for hLOX1 is mouse Anti-human LOX1 (clone 15C4, Becton Dickinson).

### Machine learning selection of ROBO2 antibodies

To predict Fab clone enrichment in later rounds of selection, a regularized logistic regression model was trained on Fab sequences from MACS and FACS1 of a ROBO2 selection campaign, where positive examples were those Fabs whose read counts had increased in FACS1 relative to MACS1. The LogisticRegression model from scikit-learn (version 1.1.3), fitted with an l1 regularization penalty strength of 1.0, was trained to predict the probability of a Fab passing FACS1. Antibody CDR3 amino acid sequences were represented as l2-normalized vectors of the counts of 1-, 2-, and 3-mers in the CDR3 sequences, and the heavy and light chain identities were represented with one-hot indicator variables. The regression parameters therefore consisted of positive or negative weights for the contribution of each k-mer and the framework identities to the likelihood of passing FACS1. A randomly selected 20% of training sequences were held out as a test set, and the enrichment of Fab sequences in FACS3 vs. MACS was used as a second test set. These hold-out sets were used to select the regularization penalty and sequence featurization schemes were selected using model performance against these hold-out sets, with the area under the receiver-operator curve (AUROC) metric. All Fabs that passed the FACS1 sort were scored by the regression model, and a subset was selected for further testing based on four criteria: 1) high logistic regression score (P(selection)>0.8); 2) sufficient read count in the FACS1 pool (>200 CPM); 3) sufficient Levenshtein distance to previously ordered sequences from FACS3 (≥5 amino acids); and 4) sufficient pairwise distance to the rest of the selected set (≥5 amino acids). From this subset, the top 29 sequences by logistic regression score were ordered.

### Immunohistochemistry staining of mouse spinal cord and brain sections for ROBO1 and ROBO2 receptors

Thirteen-day old mouse embryos (E13) were fixed by immersion in 4% paraformaldehyde in 0.12 M phosphate buffer, pH 7.4 (PFA) over night at 4°C. Samples were cryoprotected in a solution of 10% sucrose in 0.12M phosphate buffer (pH7.2), frozen in isopentane at 50°C and then cut at 20µm with a cryostat (Leica). Immunohistochemistry was performed on cryostat sections after blocking in 0.2% gelatin in PBS containing 0.5% Triton-X100 (Sigma). Sections were then incubated O/N with the following primary antibodies: goat anti-Robo1 (1:500, R&D Systems AF1749)^42^, rabbit anti-Robo2 (1:500)^42^ or IPI-ROBO1.89 and IPI-ROBO2.78 followed by 2 hours incubation in species-specific secondary antibodies directly conjugated to Cy-3 fluorophores (711-165-152 or 705-165-147 from Jackson ImmunoResearch). Sections were imaged with a fluorescent microscope (DM6000, Leica) coupled to a CoolSnapHQ camera (Roper Scientific).

## References

1. Jespers, L. S., Roberts, A., Mahler, S. M., Winter, G. & Hoogenboom, H. R. Guiding the selection of human antibodies from phage display repertoires to a single epitope of an antigen. Biotechnology (N Y) 12, 899–903 (1994).

2. Azevedo Reis Teixeira, A. et al. Drug-like antibodies with high affinity, diversity and developability directly from next-generation antibody libraries. MAbs 13, 1980942.

3. Bradbury, A. R. M., Sidhu, S., Dübel, S. & McCafferty, J. Beyond natural antibodies: the power of in vitro display technologies. Nat Biotechnol 29, 245–254 (2011).

4. Laustsen, A. H., Greiff, V., Karatt-Vellatt, A., Muyldermans, S. & Jenkins, T. P. Animal Immunization, in Vitro Display Technologies, and Machine Learning for Antibody Discovery. Trends in Biotechnology 39, 1263–1273 (2021).

5. Xu, J. L. & Davis, M. M. Diversity in the CDR3 region of V(H) is sufficient for most antibody specificities. Immunity 13, 37–45 (2000).

6. Davis, M. M. The evolutionary and structural ‘logic’ of antigen receptor diversity. Seminars in Immunology 16, 239–243 (2004).

7. Boder, E. T. & Wittrup, K. D. Yeast surface display for directed evolution of protein expression, affinity, and stability. Meth. Enzymol. 328, 430–444 (2000).

8. Lee, C. V. et al. High-affinity human antibodies from phage-displayed synthetic Fab libraries with a single framework scaffold. J Mol Biol 340, 1073–1093 (2004).

9. Mahon, C. M. et al. Comprehensive interrogation of a minimalist synthetic CDR-H3 library and its ability to generate antibodies with therapeutic potential. J Mol Biol 425, 1712–1730 (2013).

10. Sidhu, S. S. et al. Phage-displayed antibody libraries of synthetic heavy chain complementarity determining regions. J Mol Biol 338, 299–310 (2004).

11. Zhai, W. et al. Synthetic antibodies designed on natural sequence landscapes. J Mol Biol 412, 55–71 (2011).

12. Virnekäs, B. et al. Trinucleotide phosphoramidites: ideal reagents for the synthesis of mixed oligonucleotides for random mutagenesis. Nucleic Acids Res 22, 5600–5607 (1994).

13. LeProust, E. M. et al. Synthesis of high-quality libraries of long (150mer) oligonucleotides by a novel depurination controlled process. Nucleic Acids Res 38, 2522–2540 (2010).

14. Li, A., Sun, Z. & Reetz, M. T. Solid-Phase Gene Synthesis for Mutant Library Construction: The Future of Directed Evolution? Chembiochem 19, 2023–2032 (2018).

15. Olsen, T. H., Boyles, F. & Deane, C. M. Observed Antibody Space: A diverse database of cleaned, annotated, and translated unpaired and paired antibody sequences. Protein Science 31, 141–146 (2022).

16. Chen, F., Tzarum, N., Wilson, I. A. & Law, M. VH1-69 antiviral broadly neutralizing antibodies: genetics, structures, and relevance to rational vaccine design. Curr Opin Virol 34, 149–159 (2019).

17. Vargas-Madrazo, E., Lara-Ochoa, F. & Almagro, J. C. Canonical structure repertoire of the antigen-binding site of immunoglobulins suggests strong geometrical restrictions associated to the mechanism of immune recognition. J Mol Biol 254, 497–504 (1995).

18. Cobaugh, C. W., Almagro, J. C., Pogson, M., Iverson, B. & Georgiou, G. Synthetic Antibody Libraries Focused Towards Peptide Ligands. J Mol Biol 378, 622–633 (2008).

19. Valadon, P. et al. ALTHEA Gold LibrariesTM: antibody libraries for therapeutic antibody discovery. MAbs 11, 516–531 (2019).

20. Chu, X., Tian, W., Wang, Z., Zhang, J. & Zhou, R. Co-inhibition of TIGIT and PD-1/PD-L1 in Cancer Immunotherapy: Mechanisms and Clinical Trials. Mol Cancer 22, 93 (2023).

21. Akhmedov, A. et al. Lectin-like oxidized low-density lipoprotein re-ceptor-1 (LOX-1): a crucial driver of atherosclerotic cardiovascular disease. European Heart Journal 42, 1797–1807 (2021).

22. Chu, H. Y. et al. Dickkopf-1: A Promising Target for Cancer Immunotherapy. Front. Immunol. 12, (2021).

23. Pastor-Fernández, G., Mariblanca, I. R. & Navarro, M. N. Decoding IL-23 Signaling Cascade for New Therapeutic Opportunities. Cells 9, 2044 (2020).

24. Chédotal, A. Roles of axon guidance molecules in neuronal wiring in the developing spinal cord. Nature Reviews Neuroscience 1 (2019) doi:10.1038/s41583-019-0168-7.

25. Vargas, A. et al. Syncytin-2 plays an important role in the fusion of human trophoblast cells. J Mol Biol 392, 301–318 (2009).

26. Erasmus, M. F. et al. Insights into next generation sequencing guided antibody selection strategies. Sci Rep 13, 18370 (2023).

27. Morlot, C. et al. Structural insights into the Slit-Robo complex. Proceedings of the National Academy of Sciences 104, 14923–14928 (2007).

28. Green, A. G. et al. Large-scale discovery of protein interactions at residue resolution using co-evolution calculated from genomic se-quences. Nat Commun 12, 1396 (2021).

29. Cong, Q., Anishchenko, I., Ovchinnikov, S. & Baker, D. Protein interaction networks revealed by proteome coevolution. Science 365, 185–189 (2019).

30. Abramson, J. et al. Accurate structure prediction of biomolecular interactions with AlphaFold 3. Nature 630, 493–500 (2024).

31. Høie, M. H. et al. AntiFold: improved structure-based antibody design using inverse folding. Bioinformatics Advances 5, vbae202 (2025).

32. Bennett, N. R. et al. Atomically accurate de novo design of single-do-main antibodies. Preprint at 10.1101/2024.03.14.585103 (2024).

33. Erasmus, M. F. et al. AIntibody: an experimentally validated in silico antibody discovery design challenge. Nat Biotechnol 42, 1637–1642 (2024).

34. Kang, K. et al. Selection and Characterization of YKL-40-Targeting Monoclonal Antibodies from Human Synthetic Fab Phage Display Libraries. International Journal of Molecular Sciences 21, 6354 (2020).

35. Hao, Y. et al. Synthetic integrin antibodies discovered by yeast displayreveal αV subunit pairing preferences with β subunits. mAbs 16, 2365891 (2024).

36. Jo, M. H. et al. Single-molecule characterization of subtype-specific β1 integrin mechanics. Nat Commun 13, 7471 (2022).

37. Kahn, R. A. et al. Antibody characterization is critical to enhance reproducibility in biomedical research. Elife 13, e100211 (2024).

38. Jumper, J. et al. Highly accurate protein structure prediction with AlphaFold. Nature 596, 583–589 (2021).

39. Benatuil, L., Perez, J. M., Belk, J. & Hsieh, C.-M. An improved yeast transformation method for the generation of very large human antibody libraries. Protein Eng Des Sel 23, 155–159 (2010).

40. Bushnell, B., Rood, J. & Singer, E. BBMerge - Accurate paired shotgun read merging via overlap. PLoS One 12, e0185056 (2017).

41. Ghosh, A., Yang, C., Lloyd, K. & Meijers, R. High-Throughput Protein Expression Screening of Cell-Surface Protein Ectodomains. in Recombinant Protein Expression in Mammalian Cells: Methods and Protocols (ed. Hacker, D. L.) 301–316 (Springer US, New York, NY, 2024). doi:10.1007/978-1-0716-3878-1_19.

42. Dominici, C., Rappeneau, Q., Zelina, P., Fouquet, S. & Chédotal, A. Non-cell autonomous control of precerebellar neuron migration by Slit and Robo proteins. Development 145, dev150375 (2018).

