## Supplementary Tables and Figures for "High-Throughput Machine Learning-Aided Antibody Discovery for Cell Surface Antigens"

**Table S1.** Amino acid sequence liability motifs removed from CDRH3 sequences of the ARM library.

| **Rational** | **Motifs** |
|---|---|
| Oxidation | M |
| Disulfide linked aggregation | C |
| Glycosylation | NxS NxT – x not P |
| Asparagine deamidation | NG, NS, NT, NN, GNF, GNY, GNG, GNT |
| Aspartate isomerisation | DG, DS, DT, DD, DH, DK |
| Lysine glycation | KE, KD, KK, KT |
| Fragmentation | DQ |
| Polyreactivity | FF, FW, WW, GG, RR, VG, VV, YY, WxW |
| Streptavidin binding | HPQ |
| Aggregation | FHW, HYF, HWH, FDV |

**Table S2. CDRH3 length distribution for the VH1-69/VK3-15 sublibrary.** Observed CDRH3 length distribution based on Novaseq sequencing data, presented as percentage of the total sequence pool.

| CDRH3 length | Distribution (%) |
|---|---|
| 7 | 0.01 |
| 8 | 0.01 |
| 9 | 0.06 |
| 10 | 0.23 |
| 11 | 3.68 |
| 12 | 13.74 |
| 13 | 18.68 |
| 14 | 21.97 |
| 15 | 19.80 |
| 16 | 12.25 |
| 17 | 9.54 |
| 18 | 0.01 |
| 19 | 0.01 |
| 20 | 0.01 |
| 21 | 0.01 |

**Table S3.** Summary for antibodies tested and their biophysical characterization. The characterized antibodies are listed for each antigen, separated for the two library pools, namely the "flat" paratope pool consisting of light chains VK1-39/VK3-15/VK3-20 and the "concave" paratope pool consisting of light chain VK4-1. Pass/fail is reported for each antibody for polyspecificity reagent ELISA and size exclusion chromatography screens. The number of antibodies that passed both PSR and SEC with good binding kinetics for SPR (KD of less than 10 nM and an off rate lower than $10^{-2}$) and cell display titration (EC50 less than 25 nM) are also reported.

| Antigen | Light chain | Number of Abs tested | PSR pass/fail | SEC pass/fail | SPR (KD<10 nM, kd<10-2) | Cell display EC50<25 nM |
|---|---|---|---|---|---|---|
| IL23R | VK1-39/VK3-15/VK3-20 | 40 | 29/11 | 31/9 | 24 | 14 |
| LOX1 | VK1-39/VK3-15/VK3-20 | 38 | 21/17 | 26/12 | 13 | 9 |
| PDL2 | VK1-39/VK3-15/VK3-20 | 14 | 10/4 | 10/4 | 1 | 1 |
| ROBO1 | VK1-39/VK3-15/VK3-20 | 27 | 23/4 | 24/3 | 2 | 15 |
| ROBO2N | VK1-39/VK3-15/VK3-20 | 6 | 5/1 | 4/2 | 0 | 2 |
| DCC | VK1-39/VK3-15/VK3-20 | 33 | 26/7 | 21/12 | 3 | 3 |
| Syncytin | VK1-39/VK3-15/VK3-20 | 25 | 24/1 | 22/3 | 7 | 5 |
| TIGIT | VK1-39/VK3-15/VK3-20 | 28 | 22/6 | 19/9 | 3 | 7 |
| DKK | VK1-39/VK3-15/VK3-20 | 15 | 11/4 | 7/8 | 3 | 4 |
| PDL1 | VK1-39/VK3-15/VK3-20 | 27 | 21/6 | 22/5 | 7 | 15 |
| DCC | VK4-1 | 24 | 24/0 | 17/7 | 0 | 1 |
| IL23R | VK4-1 | 35 | 30/5 | 23/12 | 21 | 5 |
| LOX1 | VK4-1 | 17 | 15/2 | 10/7 | 6 | 4 |
| PDL2 | VK4-1 | 7 | 6/1 | 6/1 | 0 | 3 |
| ROBO1 | VK4-1 | 21 | 21/0 | 19/2 | 0 | 8 |
| ROBO2N | VK4-1 | 7 | 7/0 | 5/2 | 2 | 3 |
| Syncytin | VK4-1 | 36 | 36/0 | 23/13 | 11 | 12 |
| TIGIT | VK4-1 | 6 | 5/1 | 5/1 | 0 | 3 |
| DKK | VK4-1 | 7 | 7/0 | 2/5 | 0 | 0 |
| PDL1 | VK4-1 | 11 | 11/0 | 5/6 | 0 | 4 |
| Total | 20 | 424 | 354/70 | 301/123 | 103 | 118 |

**Table S4.** NGS enrichment frequencies and biophysical parameters for the most populous ARM cluster at the end of sorting for each antigen using the flat paratope (upper table) and concave epitope (lower table) sublibraries. The ARM components (heavy chain (HC), light chain (LC) and CDRH3 sequence) are reported along with relative frequency of the most abundant clone at each FACS round, as well as the biophysical characteristics obtained for these antibodies. The results of the polyspecificity reagent ELISA and size exclusion chromatography screens are reported as Pass or Fail for each antibody. Surface plasmon resonance parameters as well as the EC50 from the cell display antibody titration experiment are also reported. Note that the most abundant antibodies for human PD-L2 did not bind the target, although lower ranking antibodies did. The human DCC campaign using the VH1-69/VK4-1 sublibrary also resulted in a most dominant clone not binding the target, although lower ranked ARMs did result in productive antibodies.

Flat paratopes

| Antigen | HC | LC | CDRH3 | FACS1 | FACS2 | FACS3 | PSR | SEC | KD (nM) | ka (1/Ms) | kdis (1/s) | EC50 (nM) |
|---|---|---|---|---|---|---|---|---|---|---|---|---|
| hROBO1 | VH1-69 | VK3-15 | ARGTWIFDY | 0.0008 | 0.04 | 0.39 | Pass | Pass | NA | NA | NA | NA |
| hROBO2N | VH1-69 | VK3-15 | ARGTWIFDY | 0.003 | 0.08 | 0.95 | Pass | Pass | 22.40 | 9.40E+05 | 2.04E-02 | 0.72 |
| hIL23R_hFc | VH1-69 | VK3-15 | ARHLGSRYSSHGFDY | 0.0009 | 0.02 | 0.1 | Pass | Pass | 0.37 | 6.77E+05 | 2.54E-04 | 28.44 |
| hDCC-fn4fn6L | VH1-69 | VK3-15 | ARSWRAVDYTLFDY | 0.0004 | 0.01 | 0.21 | Pass | Fail | 29.73 | 1.84E+06 | 5.25E-02 | 37.30 |
| hTIGIT | VH1-69 | VK1-39 | ARDPRGWYGRGYAFDP | 0.01 | 0.1 | 0.74 | Pass | Fail | 12.10 | 4.00E+04 | 4.84E-03 | 2.96 |
| mDKK1 | VH1-69 | VK3-15 | ARHRYIWYRGFDY | 0.002 | 0.12 | 0.36 | Fail | Fail | 4.19 | 1.70E+05 | 7.22E-04 | 0.98 |
| hLOX1 | VH1-69 | VK1-39 | ARGQYVWNDPYFDY | 0.001 | 0.02 | 0.18 | Pass | Pass | 1.17 | 9.99E+05 | 1.17E-03 | 8.59 |
| hSyncytin2 | VH1-69 | VK3-15 | ARSGDIQWPELFDY | 0.004 | 0.1 | 0.26 | Pass | Pass | 2.36 | 1.23E+05 | 2.90E-04 | 36.13 |
| PD1-L1 | VH1-69 | VK3-20 | ARAPQYGLGYSAYFDI | 0.0001 | 0.02 | 0.2 | Pass | Pass | 3.22 | 1.30E+06 | 4.19E-03 | 0.71 |
| hPD-L2 | VH1-69 | VK3-20 | ARTPRGWYGFDY | 0.0006 | 0.05 | 0.18 | Pass | Pass | Fail | Fail | Fail | Fail |

Concave paratopes

| Antigen | HC | LC | CDRH3 | FACS1 | FACS2 | FACS3 | PSR | SEC | KD (nM) | ka (1/Ms) | kdis (1/s) | EC50 (nM) |
|---|---|---|---|---|---|---|---|---|---|---|---|---|
| hROBO1 | VH1-69 | VK4-1 | ARSERLGYWHFDY | 0.002 | 0.08 | 0.23 | Pass | Pass | NA | NA | NA | NA |
| hROBO2N | VH1-69 | VK4-1 | ARSERLGYWHFDY | 0.004 | 0.14 | 0.54 | Pass | Pass | 48.24 | 4.61E+04 | 2.23E-03 | 0.53 |
| IL-23R_hFc | VH1-69 | VK4-1 | ARGEYWYEYRTFDY | 0.002 | 0.03 | 0.16 | Pass | Pass | 1.10 | 3.57E+05 | 3.76E-04 | Fail |
| hDCC_fn4fn6L | VH1-69 | VK4-1 | ARSVQPLYPWPFDY | 0.002 | 0.02 | 0.19 | Pass | Fail | Fail | Fail | Fail | Fail |
| hTIGIT | VH1-69 | VK4-1 | ARHKGSGLVQRAVFDY | 0.02 | 0.22 | 0.94 | Pass | Pass | 87.79 | 6.68E+05 | 5.86E-02 | 2.19 |
| hDKK1 | VH1-69 | VK4-1 | AREGYGDPYAFDY | 0.003 | 0.13 | 0.62 | Pass | Fail | 16.41 | 8.36E+04 | 1.38E-03 | 73.18 |
| hLOX-1 | VH1-69 | VK4-1 | ARTAGYRWRPFDY | 0.0003 | 0.02 | 0.31 | Pass | Pass | 2.76 | 5.30E+05 | 1.47E-03 | 0.99 |
| hSyncytin2 | VH1-69 | VK4-1 | ARSGWFDEVAFDY | 0.003 | 0.02 | 0.09 | Pass | Pass | 5.23 | 5.41E+04 | 2.82E-04 | 0.98 |
| hPD1-L1 | VH1-69 | VK4-1 | ARANFSAAFDRAFDP | 0.0006 | 0.04 | 0.42 | Pass | Fail | 9.34 | 4.80E+06 | 4.48E-02 | 0.26 |
| hPD-L2 | VH1-69 | VK4-1 | ARAGLPAPGEVKVFDY | 0.003 | 0.1 | 0.55 | Pass | Pass | Fail | Fail | Fail | Fail |

**Table S5.** ML-based ranking of ROBO2N ARM clones that were lost during cell sorting, using logistic regression on k-mer selections between the MACS and FACS1 rounds by LR score. Antibodies that showed positive binding in SPR and cell display are colored green. The table combines antibody information (CDRH3 sequence, heavy chain (HC), light chain (LC) with deep sequencing data (frequency per ARM clone for each FACS round) and biophysical data, including pass/fail for the ELISA polyspecificity reagent and size exclusion chromatography screen, surface plasmon resonance parameters (Kd and kOff) and antibody potency as derived from cell display of the target antigen (EC50). The LR score is the logistic regression score based on 1, 2 and 3 k-mers that are enriched in the FACS1 round compared to the preceding MACS round.

| Antibody name | HCDR3 | HC | LC | FACS1 | FACS2 | FACS3 | SPR Kd | SPR kOff | Cell display EC50 (nM) | LR_score |
|---|---|---|---|---|---|---|---|---|---|---|
| ROBO2N_ML_Ab_001 | ARSKYVYWGDAFDY | VH1-69 | VK3-15 | 4.16E-04 | 1.66E-03 | 0 | 9.96E-08 | 1.3E-03 | 3.5 | 0.99 |
| ROBO2N_ML_Ab_002 | ARHGTIRPLTWGDAFDI | VH1-69 | VK3-15 | 7.43E-04 | 0 | 0 | 1.34E-07 | 1.7E-03 | 9.8 | 0.98 |
| ROBO2N_ML_Ab_003 | ARGRPHRWTAWGDAFDI | VH1-69 | VK3-15 | 1.59E-03 | 0 | 0 | 9.42E-08 | 8.3E-04 | 1.0 | 0.98 |
| ROBO2N_ML_Ab_004 | ARAEPARVSYWGDAFDI | VH1-69 | VK3-15 | 1.26E-05 | 1.47E-03 | 0 | 1.94E-07 | 2.6E-03 | 6.2 | 0.97 |
| ROBO2N_ML_Ab_005 | ARSWIQLYAFDY | VH1-69 | VK3-15 | 4.79E-04 | 9.48E-05 | 0 | 1.68E-07 | 1.8E-03 | 9.8 | 0.94 |
| ROBO2N_ML_Ab_006 | ARSQSSQYSGWGDAFDI | VH1-69 | VK3-15 | 6.05E-04 | 0 | 0 | 2.37E-07 | 2.8E-03 | Fail | 0.92 |
| ROBO2N_ML_Ab_007 | ARHQRGQYVAWGNAFDY | VH1-69 | VK3-15 | 1.12E-03 | 2.84E-04 | 0 | 1.42E-07 | 1.5E-03 | 17.6 | 0.91 |
| ROBO2N_ML_Ab_008 | ARHTPTHYALWGDAFDY | VH1-69 | VK3-15 | 4.28E-04 | 0 | 0 | 1.45E-07 | 1.3E-03 | 125.0 | 0.91 |
| ROBO2N_ML_Ab_009 | ARNAEPNISGSAFDI | VH1-69 | VK3-15 | 2.90E-04 | 0 | 0 | 1.61E-07 | 3.2E-03 | 3.5 | 0.89 |
| ROBO2N_ML_Ab_010 | ARVHSIRYWLFDY | VH1-69 | VK3-15 | 6.05E-04 | 0 | 0 | Fail | Fail | Fail | 0.88 |
| ROBO2N_ML_Ab_011 | ARSIDIATNEGYFDP | VH1-69 | VK3-20 | 3.78E-04 | 0 | 0 | Fail | Fail | Fail | 0.87 |
| ROBO2N_ML_Ab_012 | ARGVEPYLAGSGFDI | VH1-69 | VK3-15 | 1.21E-03 | 5.03E-03 | 0 | 1.14E-07 | 1.9E-03 | 1.5 | 0.87 |
| ROBO2N_ML_Ab_013 | ARGAKSYNQIGFDY | VH1-69 | VK1-39 | 7.56E-04 | 0 | 0 | Fail | Fail | Fail | 0.86 |
| ROBO2N_ML_Ab_014 | ARGYEAWYSGSAFDP | VH1-69 | VK3-15 | 1.51E-03 | 5.38E-03 | 0 | 6.28E-08 | 1.2E-03 | 4.4 | 0.86 |
| ROBO2N_ML_Ab_015 | ARGPLTWTASETDAFDY | VH1-69 | VK1-39 | 5.29E-04 | 0 | 0 | Fail | Fail | Fail | 0.85 |
| ROBO2N_ML_Ab_016 | ARVRWVLYAGFDY | VH1-69 | VK3-15 | 3.28E-04 | 0 | 0 | Fail | Fail | Fail | 0.85 |
| ROBO2N_ML_Ab_017 | ARSPWLRNAYFDY | VH1-69 | VK3-20 | 1.89E-04 | 0 | 0 | Fail | Fail | Fail | 0.85 |
| ROBO2N_ML_Ab_018 | ARENWNYAVYFPFDY | VH1-69 | VK3-20 | 1.76E-04 | 0 | 0 | Fail | Fail | Fail | 0.85 |
| ROBO2N_ML_Ab_019 | ARIYGVTYVWYFDP | VH1-69 | VK3-15 | 6.05E-04 | 0 | 0 | Fail | Fail | Fail | 0.85 |
| ROBO2N_ML_Ab_020 | ARAILWLTTALAGFDY | VH1-69 | VK3-15 | 1.26E-05 | 0 | 0 | Fail | Fail | Fail | 0.84 |
| ROBO2N_ML_Ab_021 | ARDRPDLWLPRGFDY | VH1-69 | VK3-20 | 7.18E-04 | 0 | 0 | Fail | Fail | Fail | 0.84 |
| ROBO2N_ML_Ab_022 | ARVYGSYPRGIFDY | VH1-69 | VK3-15 | 8.06E-04 | 0 | 0 | Fail | Fail | Fail | 0.84 |
| ROBO2N_ML_Ab_023 | ARVSSFDAYPGFDY | VH1-69 | VK3-15 | 2.90E-04 | 0 | 0 | Fail | Fail | Fail | 0.84 |
| ROBO2N_ML_Ab_024 | ARNAERWLAGSYFDP | VH1-69 | VK3-15 | 1.17E-03 | 9.15E-03 | 0 | 7.96E-08 | 1.2E-03 | 7.8 | 0.84 |
| ROBO2N_ML_Ab_025 | ARDEVEWGVQLYNAFDI | VH1-69 | VK1-39 | 1.26E-05 | 0 | 0 | Fail | Fail | Fail | 0.84 |
| ROBO2N_ML_Ab_026 | ARNPSRESGWGDPFDY | VH1-69 | VK3-20 | 2.14E-04 | 0 | 0 | Fail | Fail | Fail | 0.83 |
| ROBO2N_ML_Ab_027 | ARDLHIGRLANYFDP | VH1-69 | VK3-15 | 2.52E-04 | 0 | 0 | Fail | Fail | Fail | 0.83 |
| ROBO2N_ML_Ab_028 | ARAIGHFERQGFDY | VH1-69 | VK1-39 | 1.26E-05 | 0 | 0 | Fail | Fail | Fail | 0.83 |
| ROBO2N_ML_Ab_029 | ARVDESPLTYFDY | VH1-69 | VK1-39 | 2.65E-04 | 0 | 0 | Fail | Fail | Fail | 0.83 |

**Table S6.** ML-based ranking of PDL2 ARM clones that were lost during cell sorting, using logistic regression on k-mer selections between the MACS and FACS1 rounds by LR score. Antibodies that showed positive binding in SPR and cell display are colored green. The table combines antibody information (CDRH3 sequence, heavy chain (HC), light chain (LC) with deep sequencing data (frequency per ARM clone for each FACS round) and biophysical data, including pass/fail for the ELISA polyspecificity reagent and size exclusion chromatography screen, surface plasmon resonance parameters (Kd and kOff) and antibody potency as derived from cell display of the target antigen (EC50). The LR score is the logistic regression score based on 1, 2 and 3 k-mers that are enriched in the FACS1 round compared to the preceding MACS round. The two experimentally obtained PDL2 antibodies from the same campaign are added in red for comparison.

| | | | | | | | | | | |
|---|---|---|---|---|---|---|---|---|---|---|
| hPD-L2_ML1 | CARSHRRSVDFDY | VH1-69 | VK1-39 | 4.16E-04 | 0 | 0 | 3.00E-08 | 2.81E-02 | 1.58 | 0.91 |
| hPD-L2_ML2 | CARPHDYREWLFDY | VH1-69 | VK3-15 | 6.69E-04 | 0 | 0 | 1.95E-07 | 8.60E-02 | 2.53 | 0.90 |
| hPD-L2_ML3 | CARGASTSNYGFDY | VH1-69 | VK3-15 | 3.72E-04 | 0 | 0 | 1.78E-07 | 1.17E-01 | 3.91 | 0.89 |
| hPD-L2_ML4 | CARVARGRYNLIFDY | VH1-69 | VK1-39 | 2.23E-04 | 0 | 0 | Fail | Fail | Fail | 0.89 |
| hPD-L2_ML5 | CARSYRRLPAFDY | VH1-69 | VK3-20 | 1.13E-03 | 0 | 0 | Fail | Fail | Fail | 0.88 |
| hPD-L2_ML6 | CARSYRLYIGFDY | VH1-69 | VK3-15 | 1.13E-03 | 0 | 0 | 9.97E-08 | 2.68E-02 | 4.20 | 0.87 |
| hPD-L2_ML7 | CARGRYRIYIVSQPFDY | VH1-69 | VK1-39 | 2.83E-04 | 0 | 0 | Fail | Fail | Fail | 0.87 |
| hPD-L2_ML8 | CARWADYRDPLFDY | VH1-69 | VK3-20 | 3.42E-04 | 0 | 0 | Fail | Fail | Fail | 0.87 |
| hPD-L2_ML9 | CARSYRQEVLFDY | VH1-69 | VK3-15 | 2.53E-04 | 0 | 0 | 3.67E-07 | 1.58E-01 | 22.49 | 0.86 |
| hPD-L2_ML10 | CARVARGVYDLLFDY | VH1-69 | VK1-39 | 3.42E-04 | 0 | 0 | Fail | Fail | Fail | 0.86 |
| hPD-L2_ML11 | CARVPRGAVSPWPFDY | VH1-69 | VK3-20 | 6.54E-04 | 0 | 0 | Fail | Fail | Fail | 0.86 |
| hPD-L2_ML12 | CARNLGKNDYFQVYFDY | VH1-69 | VK1-39 | 3.12E-04 | 0 | 0 | 6.88E-08 | 2.69E-02 | 4.54 | 0.86 |
| hPD-L2_ML13 | CARVARTAERLDFDY | VH1-69 | VK3-20 | 2.23E-04 | 0 | 0 | Fail | Fail | Fail | 0.86 |
| hPD-L2_ML14 | CARSPRQSTVFDY | VH1-69 | VK3-15 | 2.83E-04 | 0 | 0 | 2.94E-08 | 3.28E-02 | 0.51 | 0.86 |
| hPD-L2_ML15 | CARNYRQDLHFDY | VH1-69 | VK1-39 | 5.35E-04 | 0 | 0 | 3.14E-07 | 1.25E-01 | 29.79 | 0.86 |
| hPD-L2_ML16 | CARGADYRDVSFDY | VH1-69 | VK1-39 | 1.09E-03 | 0 | 0 | 1.90E-08 | 1.28E-02 | 1.11 | 0.85 |
| hPD-L2_ML17 | CARSHRSVWLFDY | VH1-69 | VK3-15 | 6.39E-04 | 0 | 0 | 1.83E-06 | 1.19E-01 | 81.76 | 0.85 |
| hPD-L2_ML18 | CARGEDYRAVDFDY | VH1-69 | VK1-39 | 4.61E-04 | 0 | 0 | 6.77E-08 | 1.21E-02 | 5.14 | 0.85 |
| hPD-L2_ML19 | CARSYRLYGSLFDY | VH1-69 | VK1-39 | 2.53E-04 | 0 | 0 | Fail | Fail | Fail | 0.84 |
| hPD-L2_ML20 | CARPSDYRTVDFDY | VH1-69 | VK1-39 | 5.21E-04 | 0 | 0 | 2.61E-08 | 2.50E-02 | 2.33 | 0.84 |
| hPD-L2_ML21 | CARSPRQRVDFDY | VH1-69 | VK1-39 | 8.18E-04 | 0 | 0 | 3.23E-08 | 2.37E-02 | 1.05 | 0.84 |
| hPD-L2_ML22 | CARLVKRNIGAFDY | VH1-69 | VK3-15 | 5.35E-04 | 0 | 0 | 1.13E-07 | 5.54E-02 | 1.45 | 0.84 |
| hPD-L2_ML23 | CARSHRSSIAPYFDY | VH1-69 | VK1-39 | 2.08E-04 | 0 | 0 | Fail | Fail | Fail | 0.83 |
| hPD-L2_ML24 | CARGIRINYVSFDY | VH1-69 | VK1-39 | 3.42E-04 | 0 | 0 | 7.80E-08 | 5.41E-02 | 0.79 | 0.83 |
| hPD-L2_ML25 | CARVKRQHTQFDY | VH1-69 | VK3-20 | 6.25E-04 | 0 | 0 | Fail | Fail | Fail | 0.82 |
| hPD-L2_ML26 | CARVPRQVSNEWSAFDY | VH1-69 | VK1-39 | 2.53E-04 | 0 | 0 | Fail | Fail | Fail | 0.82 |
| hPD-L2_ML27 | CARSYRTGWLFDY | VH1-69 | VK3-15 | 1.32E-03 | 0 | 0 | 3.40E-07 | 1.05E-01 | 16.78 | 0.82 |
| hPD-L2_ML28 | CARLPRQQSEFDY | VH1-69 | VK3-20 | 1.50E-03 | 0 | 0 | Fail | Fail | Fail | 0.82 |
| hPD-L2_ML29 | CARVPRWSSYIPAGFDY | VH1-69 | VK3-20 | 3.57E-04 | 0 | 0 | Fail | Fail | Fail | 0.81 |
| hPD-L2_ML30 | CARGSRIYDVSFDY | VH1-69 | VK3-15 | 2.23E-04 | 0 | 0 | Fail | Fail | Fail | 0.81 |
| hPD-L2_ML31 | CARELVDVDYRLGSFDI | VH1-69 | VK3-15 | 2.53E-04 | 0 | 0 | Fail | Fail | Fail | 0.81 |
| hPD-L2_ML32 | CARGSHRARVSFDY | VH1-69 | VK1-39 | 2.38E-04 | 0 | 0 | 6.50E-08 | 6.05E-02 | 0.60 | 0.80 |
| hPD-L2_ML33 | CARTVKGQWLFDY | VH1-69 | VK3-15 | 5.21E-04 | 0 | 0 | Fail | Fail | Fail | 0.80 |
| hPD-L2_exp1 | CARSLGRSYTFQVYFDY | VH1-69 | VK1-39 | 3.27E-04 | 2.52E-04 | 2.19E-02 | 1.61E-01 | 1.53E-03 | 0.99 | NA |
| hPD-L2_exp2 | CARTSIQYNRWLAEGFDP | VH1-69 | VK3-15 | 5.38E-04 | 6.28E-03 | 1.36E-02 | 4.20E-08 | 5.59E-03 | 2.66 | NA |

**Table S7.** Oligos used in NGS.

| Oligonucleotide Name | 5'-3' Primer Sequence |
|---|---|
| NGS_FA | GTCTCGTGGGCTCGGAGATGTGTATAAGAGACAGCCAGGTAAAGGTTTGGAATGG |
| NGS_Rd | TCGTCGGCAGCGTCAGATGTGTATAAGAGACAGTGGAGGAGGGTGCCAGAG |
| NGS_Re | TCGTCGGCAGCGTCAGATGTGTATAAGAGACAGHHTGGAGGAGGGTGCCAGAG |
| NGS_Rf | TCGTCGGCAGCGTCAGATGTGTATAAGAGACAGHHHHTGGAGGAGGGTGCCAGAG |

**TableS8.** Description of antigen constructs

| Antigen | Uniprot ID | Residues | Signal Sequence | Signal Sequence name |
|---|---|---|---|---|
| PD-L1 | Q9NZQ7 | 18 – 238 | MDWTWRILFLVAAATGAHS | hIgVH [1] |
| PD-L2 | Q9BQ51 | 20 – 220 | MDWTWRILFLVAAATGAHS | hIgVH [1] |
| TIGIT | Q495A1 | 22 – 141 | MRVPAQLLGLLLLWFPGSRC | hIgKappa [2] |
| LOX1 | P78380 | 57 – 273 | MRVPAQLLGLLLLWFPGSRC | hIgKappa [2] |
| DKK1 | O54908 | 32 – 272 | MDWTWRILFLVAAATGAHS | hIgVH [1] |
| IL23 | Q5VWK5 | 24 – 355 | MPLLLLLPLLWAGALA | CD33 [2] |
| DCC | P43146 | 721 – 1043 | MRVPAQLLGLLLLWFPGSRC | hIgKappa [2] |
| Syncytin-2 | P60508 | 16 – 467 | MRVPAQLLGLLLLWFPGSRC | hIgKappa [2] |
| ROBO1 | Q9Y6N7 | 26 – 897 | MDWTWRILFLVAAATGAHS | hIgVH [1] |
| ROBO 2N | Q9HCK4 | 21 – 129 | MRVPAQLLGLLLLWFPGSRC | hIgKappa [2] |

1. Haryadi et al Plos One 2015[1]
2. Güler-Gane et al PLoS One 2016[2]

**Table S9.** Cell display constructs for the 10 cell surface antigens

| | Display construct | UniProt | Sequence | Signal sequence | SP name | Manufacturer |
|---|---|---|---|---|---|---|
| IL23R | IL23R_HUMAN | Q5VWK5 | 24-355 | MPLLLLLPLLWAGALA | hCD33_SP | Synbio |
| | IL23R_MOUSE | Q8K4B4 | 24-374 | MPLLLLLPLLWAGALA | hCD33_SP | Synbio |
| ROBO1 | ROBO1_HUMAN | Q9Y6N7 | 26-897 | MPLLLLLPLLWAGALA | hCD33_SP | GenScript |
| | ROBO1_MOUSE | O89026 | 21-858 | MPLLLLLPLLWAGALA | hCD33_SP | GenScript |
| ROBO2 | ROBO2_HUMAN | Q9HCK4 | 22-859 | MPLLLLLPLLWAGALA | hCD33_SP | GenScript |
| | ROBO2_MOUSE | Q7TPD3 | 23-864 | MPLLLLLPLLWAGALA | hCD33_SP | GenScript |
| PDL1 | PDL1_HUMAN | Q9NZQ7 | 19-238 | MTRLTVLALLAGLLASSRA | hAZU1_SP | TWIST |
| | PDL1_MOUSE | Q9EP73 | 20-238 | MTRLTVLALLAGLLASSRA | hAZU1_SP | TWIST |
| PDL2 | PDL2_HUMAN | Q9BQ51 | 20-220 | MTRLTVLALLAGLLASSRA | hAZU1_SP | TWIST |
| | PDL2_MOUSE | Q9WUL5 | 20-220 | MTRLTVLALLAGLLASSRA | hAZU1_SP | TWIST |
| TIGIT | TIGIT_HUMAN | Q495A1 | 22-141 | MPLLLLLPLLWAGALA | hCD33_SP | Synbio |
| | TIGIT_MOUSE | P86176 | 18-147 | MPLLLLLPLLWAGALA | hCD33_SP | Synbio |
| DKK1 | DKK1_HUMAN | O94907 | 31-266 | MPLLLLLPLLWAGALA | hCD33_SP | Synbio |
| | DKK1_MOUSE | O54908 | 32-272 | MPLLLLLPLLWAGALA | hCD33_SP | Synbio |
| DCC | DCC_HUMAN | P43146 | 26-1097 | MPLLLLLPLLWAGALA | hCD33_SP | Synbio |
| | DCC_MOUSE | P70211 | 26-1097 | MPLLLLLPLLWAGALA | hCD33_SP | Synbio |
| LOX1 | LOX1_HUMAN | P78380 | 58-273 | MERVQPLEENVGNAARPRFERNKLLLVASVIQGLGLLLCFTYICLHFSAL | hOX40L_leader_sequence | Synbio |

| | | | | | | |
|---|---|---|---|---|---|---|
| | LOX1_MOUSE | Q9EQ09 | 55-363 | MERVQPLEENVGNAARPRFERNKLLLVASVIQGLGLLLCFTYICLHFSAL | hOX40L_leader_sequence | Synbio |
| SYNC2 | SYNC2_FL | P60508 | 16-538 | MTRLTVLALLAGLLASSRA | hAZU1_SP | TWIST |
| | SYNC2_S2 | P60508 | 351-538 | MTRLTVLALLAGLLASSRA | hAZU1_SP | TWIST |

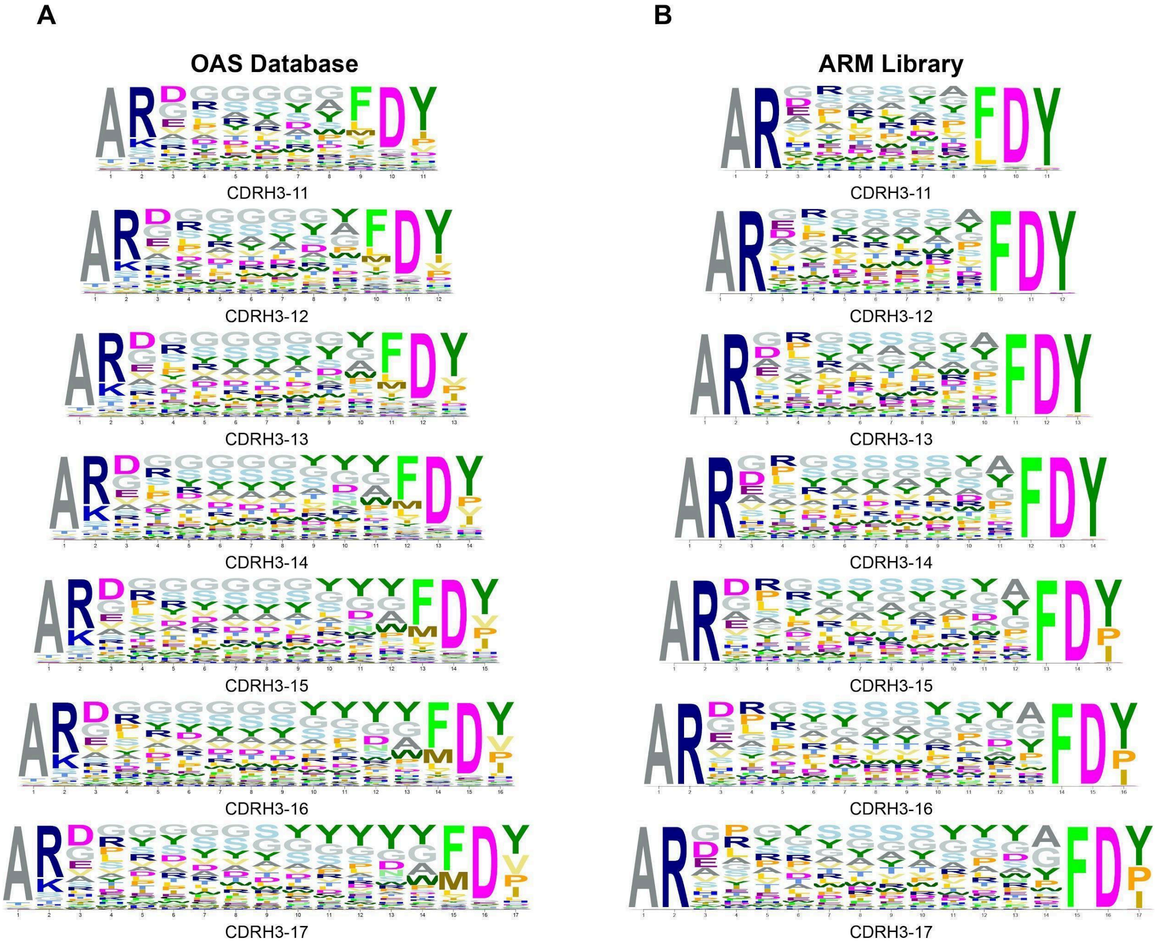


**Figure S1**. Sequence logos displaying amino acid frequencies for each CDRH3 region contributing to the antigen recognition module. Amino acid frequencies were calculated using human naïve B cell sequencing data (A) from the Observed Antibody Space (OAS) database, (B) shows amino acid frequencies obtained from the NovaSeq dataset of the VH1-69/VK3-15 sublibrary. The logos were made with Weblogo[3].

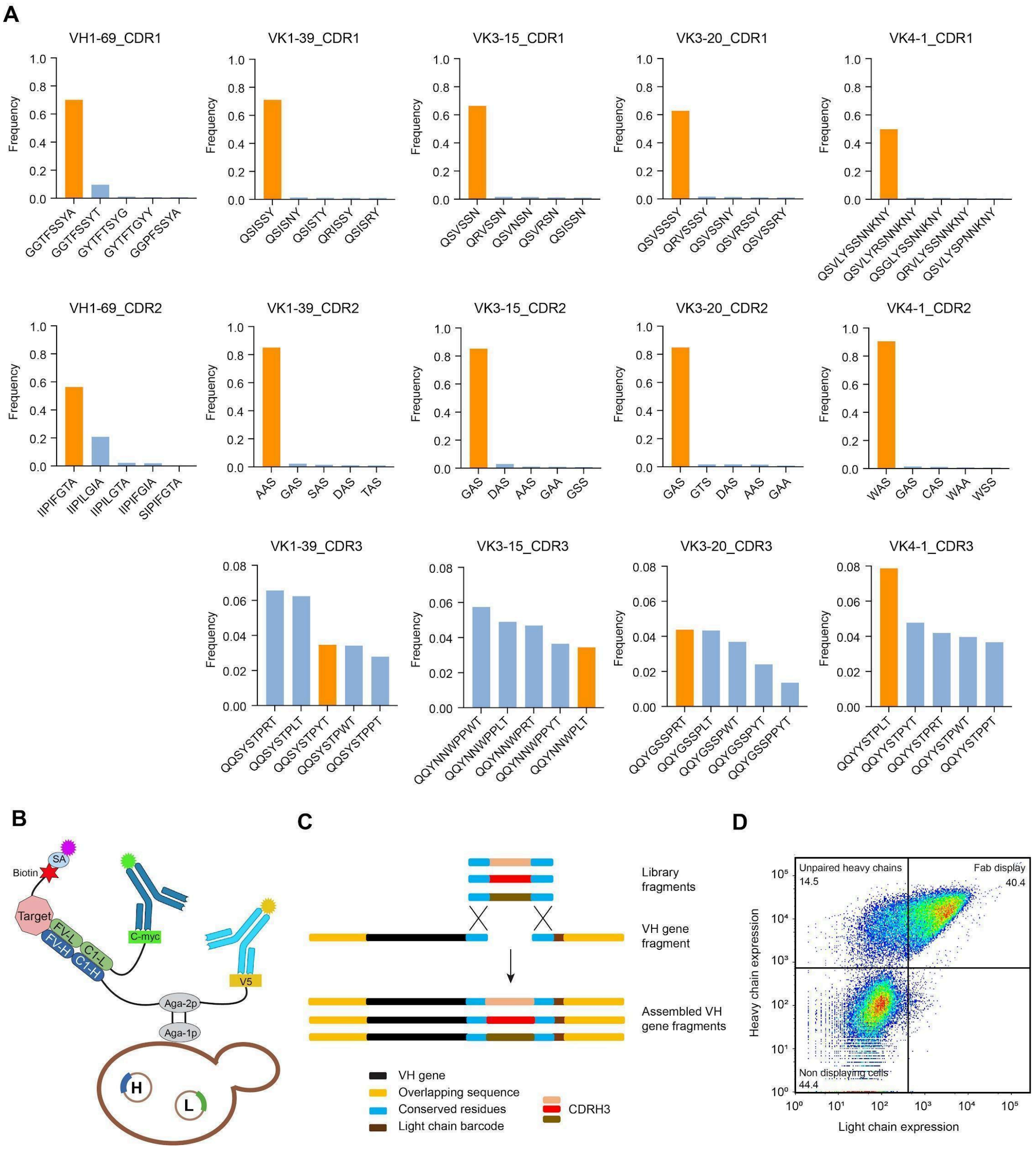


**Figure S2.** Library assembly. A) Frequencies of CDR sequences for the VH1-69 heavy chain and the light chains used in the library, as observed in the OAS database. The frequencies of the top 5 sequences for each CDR (H1, H2, L1, L2, and L3) corresponding to heavy and light chains are shown as bar graphs. The orange bars indicate the CDR sequences chosen for the library. B) Schematic representation of IPI's yeast display platform. A two-vector system is used to display Fabs on the yeast surface. The heavy chain is fused to the N-terminus of Aga2 protein, and its display is detected by staining for a V5 tag. The light chain has a C-Myc tag, and antigen binding is detected using fluorescently labeled streptavidin. C) Schematic showing library assembly, the light chain barcode region highlighted in brown. D) FACS plot showing display of the mixed VH1-69 Fab library. The top right quadrant indicates cells displaying paired Fabs, the top left indicates unpaired Fabs, and the bottom left indicates non-displaying cells.

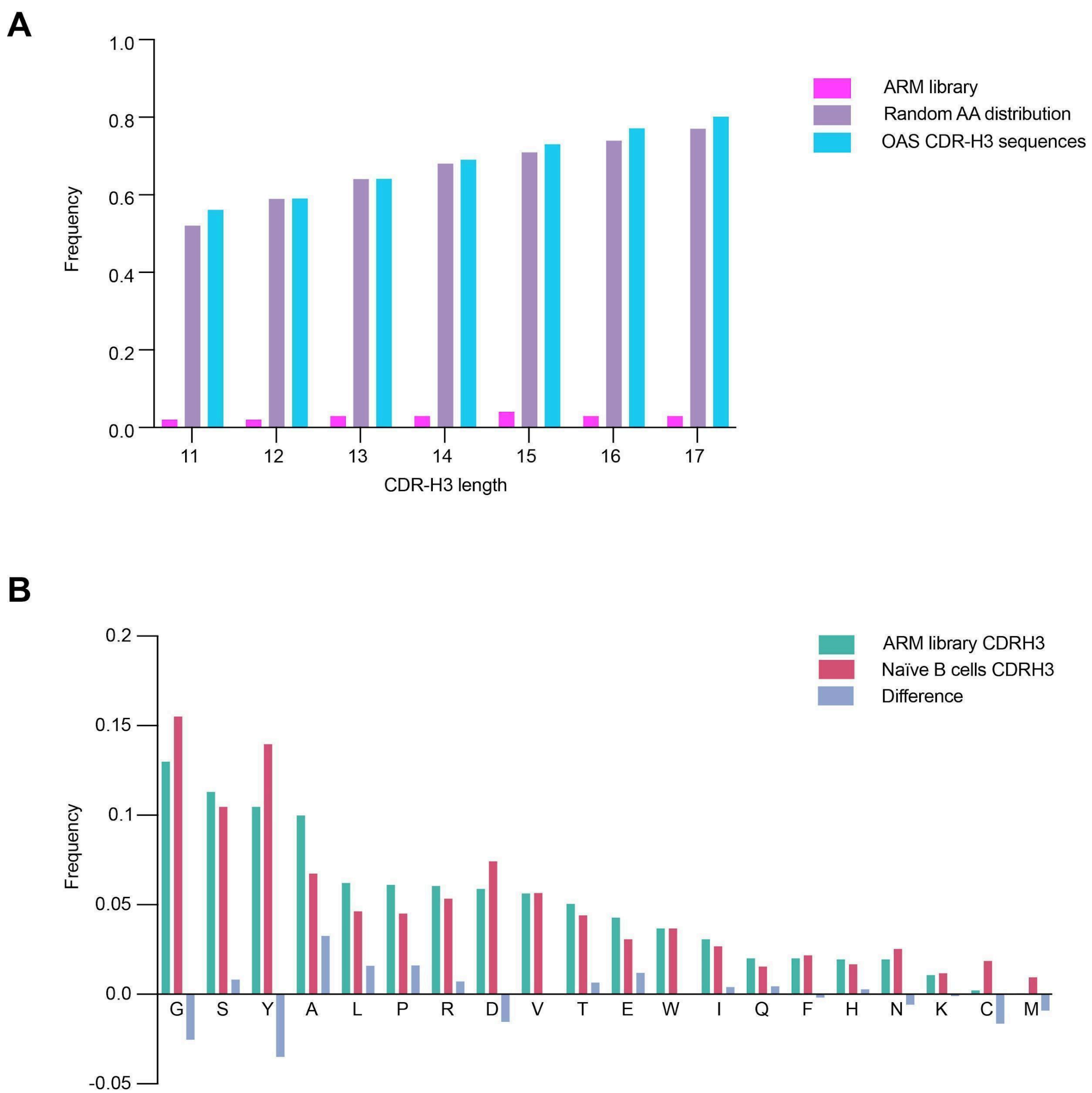


**Figure S3.** **(A)** Liability motif prevalence in the ARM library compared to a random amino acid distribution library, and naïve B-cell CDRH3 sequences from the OAS database. **(B)** Bar graph comparing the amino acid frequencies in the CDRH3 region of the library (blue) to the naïve B-cell sequencing dataset (crimson). Differences in frequency are shown in grey.

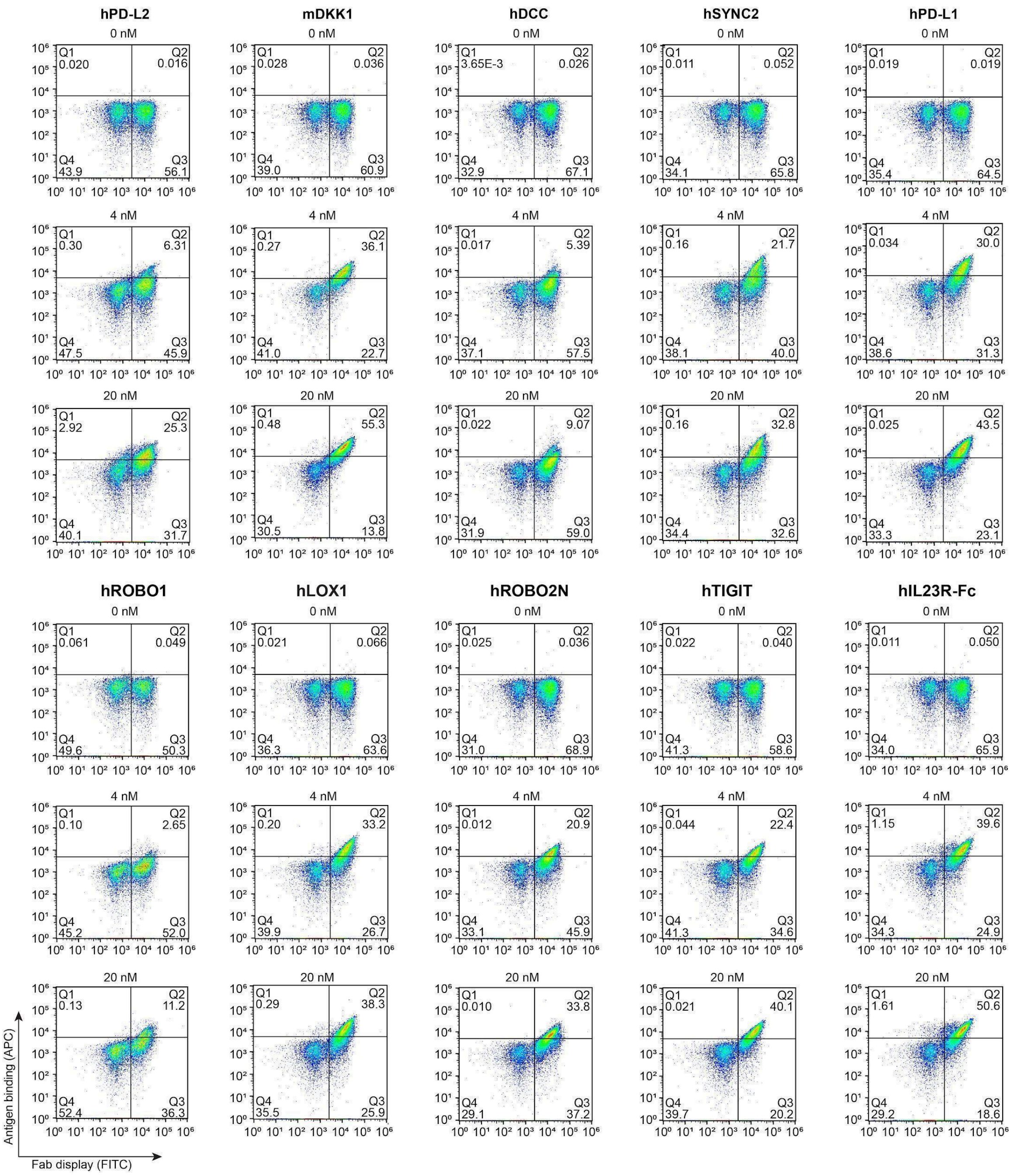


**Figure S4.** FACS plots showing yeast Fab clones binding at 0 nm, 4 nM and 20 nM antigen concentrations for the flat paratope library to 10 antigens. The top right quadrant shows Fab clones binding to Streptavidin-APC linked antigen, the bottom right shows yeast clones that display Fabs but does not bind to the antigen (stained with a FITC conjugated anti-Myc tag antibody), and the bottom left indicates non-displaying yeast cells.

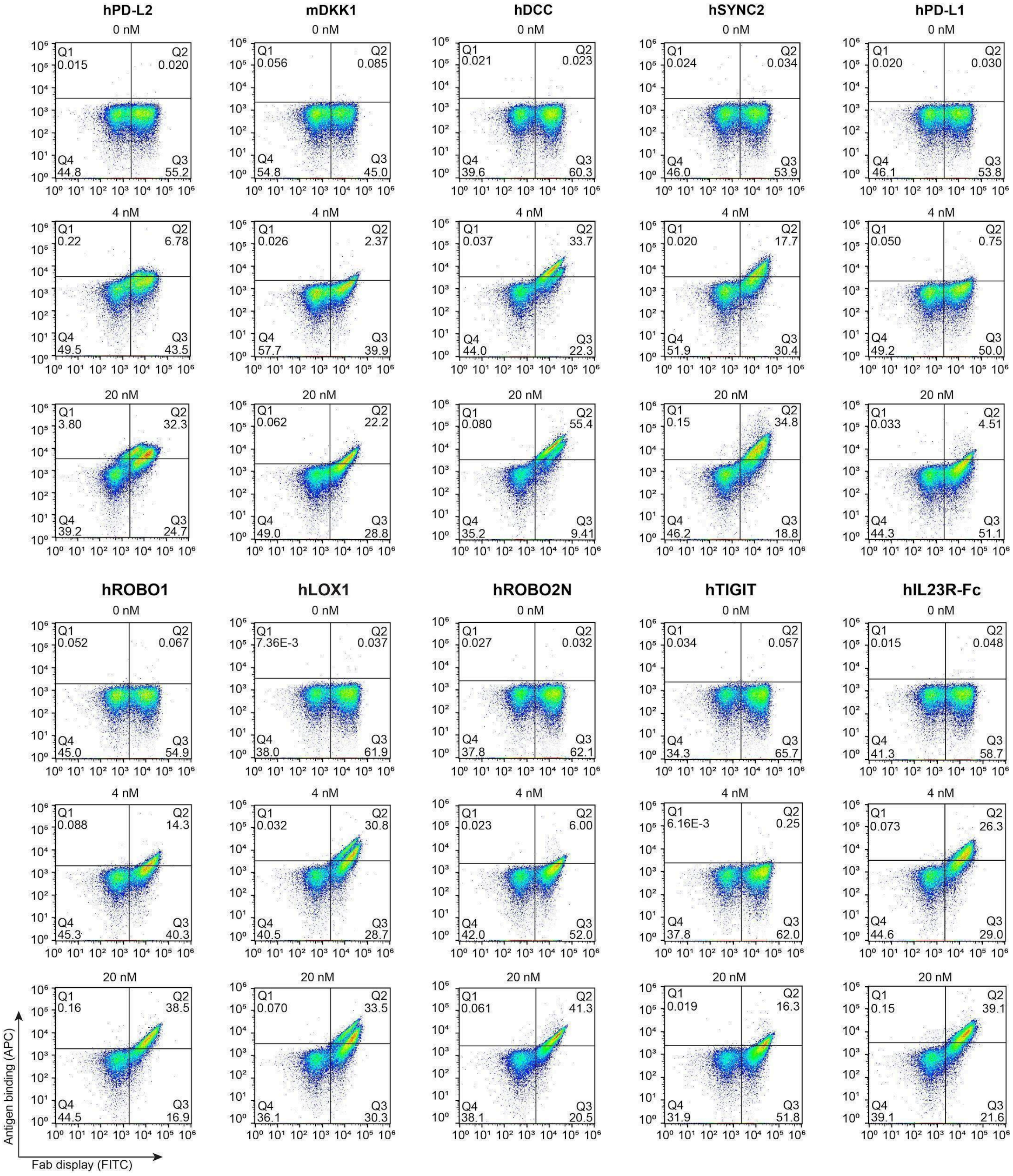


**Figure S5.** FACS plots showing yeast Fab clones binding at 0 nM, 4 nM and 20 nM antigen concentrations for the concave paratope library to 10 antigens The top right quadrant shows Fab clones binding to Streptavidin-APC linked antigen, the bottom right shows yeast clones that display Fabs but does not bind to the antigen(stained with a FITC conjugated anti-Myc tag antibody), and the bottom left indicates non-displaying yeast cells.

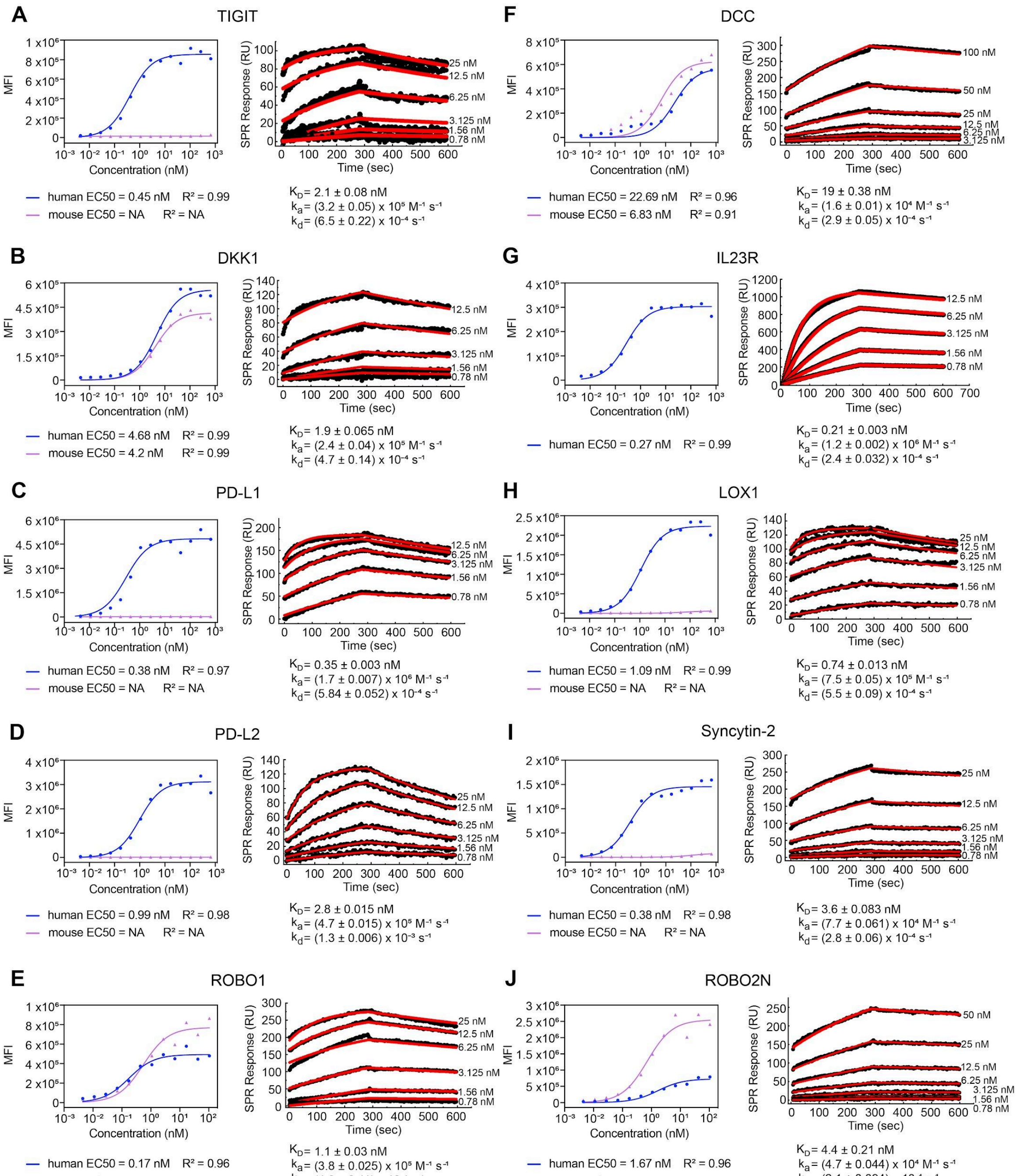

**Figure S6.** Cell display antibody titration curves and surface plasmon resonance sensograms for the best performing antibodies for TIGIT, DKK, PD-L1, PD-L2, DCC, IL23R, LOX1, Syncytin-2, ROBO1 and ROBO2 that show consistent and optimal binding kinetics. For cell display, both human (blue) and murine (red) antigen was displayed on Expi-CHO cells to verify species crossreactivity.

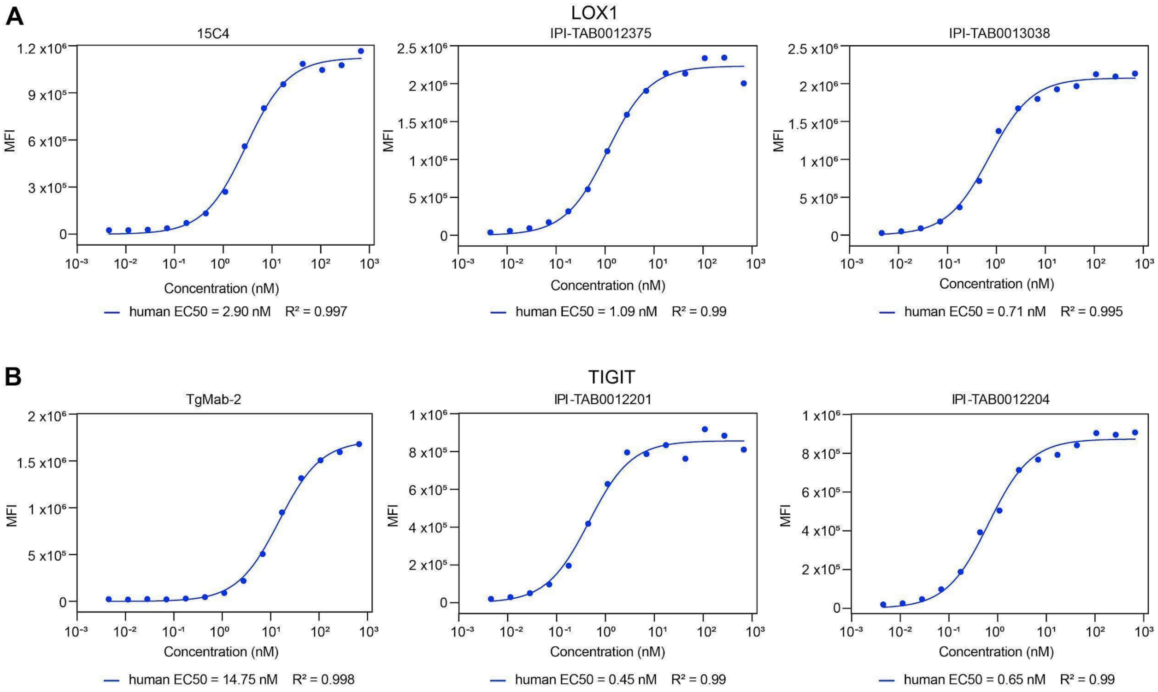


**Figure S7.** Benchmarking IPI's antibodies for flow cytometry. IPI antibodies (in human IgG1 format) targeting human LOX1 and TIGIT were compared to commercially available antibodies 15C4 (mIgG2a anti-human LOX1) and TgMab2 (mIgG1 anti-human TIGIT) from BD Biosciences. LOX1 and TIGIT full length ectodomains were displayed on CHO cell surfaces, and binding was assessed at multiple antibody concentrations by Alexa647 labeled anti-Fc secondary for the respective species. IPI antibodies had comparable (LOX1) or substantially lower (TIGIT) $EC_{50}$ values than the commercial clones.

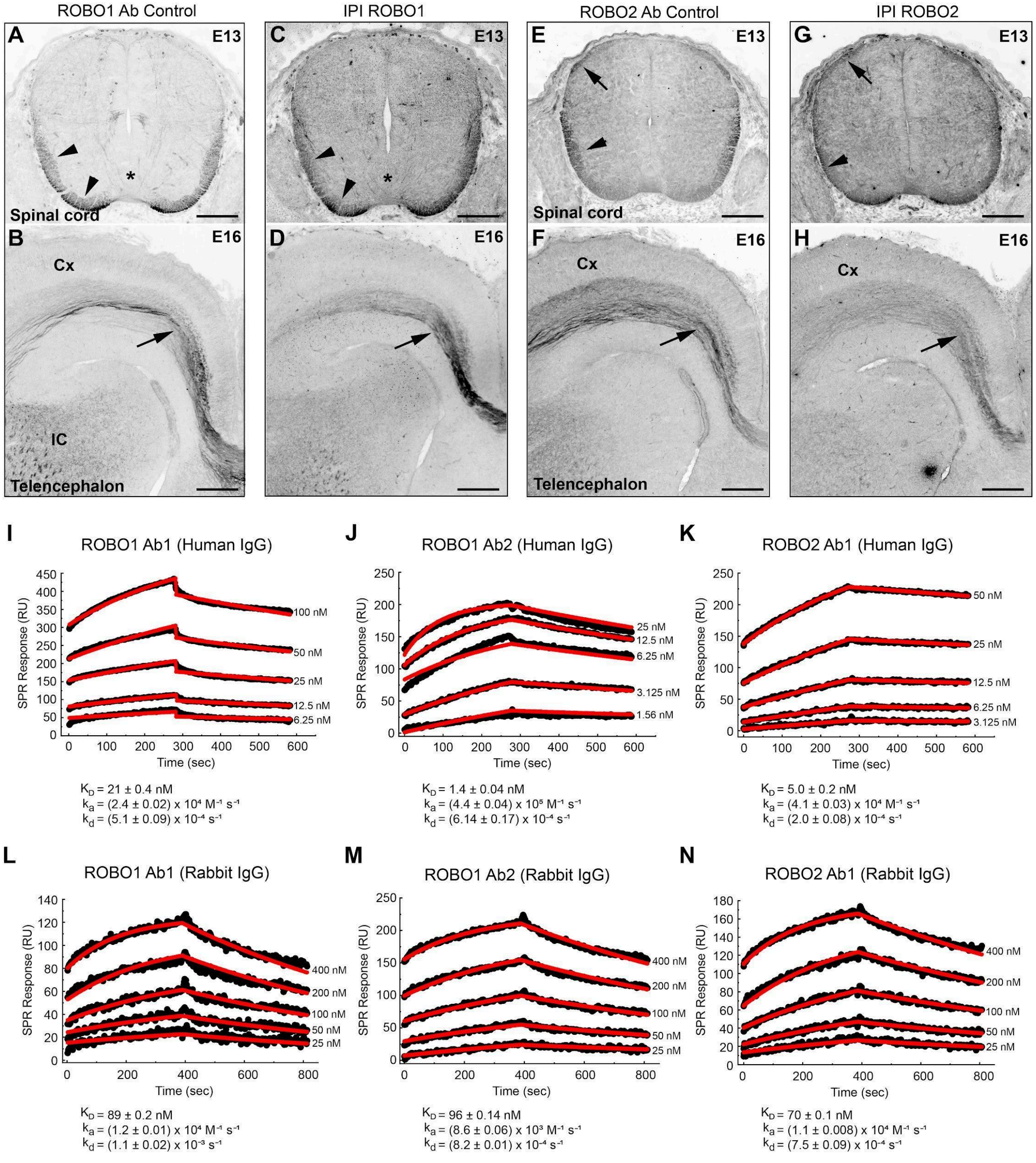


**Figure S8.** Examples of IHC staining of selected antibodies for ROBO1 and ROBO2 receptors. Cryostat sections of E13 mouse spinal cord (A, C, E, G) or E16 telencephalon (B, D, F, H) were labeled with positive control, previously validated Robo1 (A, B) or Robo2 (E, F) antibodies or with IPI human/rabbit chimera IgG ROBO1.89 (C, D) or ROBO2.78 (G, H) IPI antibodies. (A, C, E, G) at E13, post-crossing commissural axons extending under the surface of the ventral and lateral spinal cord (arrowheads) are immunoreactive for Robo1 or Robo2. Robo2 immunoreactivity is also detected in sensory axons at the dorsal root entry zone (arrow in E and G). The floor plate, or ventral midline of the spinal cord, is indicated with an asterisk (A, C). (B, D, F, H) at E16, callosal axons connecting the two sides of the neocortex are immunoreactive for both control and IPI ROBO1 (B, D) and ROBO2 (F, H) antibodies. In the striatum, axons of the internal capsule (IC) also express Robo1 (B, D). Scales bars: 150μm (A, C, E, G); 250μm (B, D, F, H). (I- N) Comparison of the SPR kinetics of the human IgG1 and human-rabbit chimera IgG antibodies for ROBO1 and ROBO2 antibodies used in the IHC experiments.

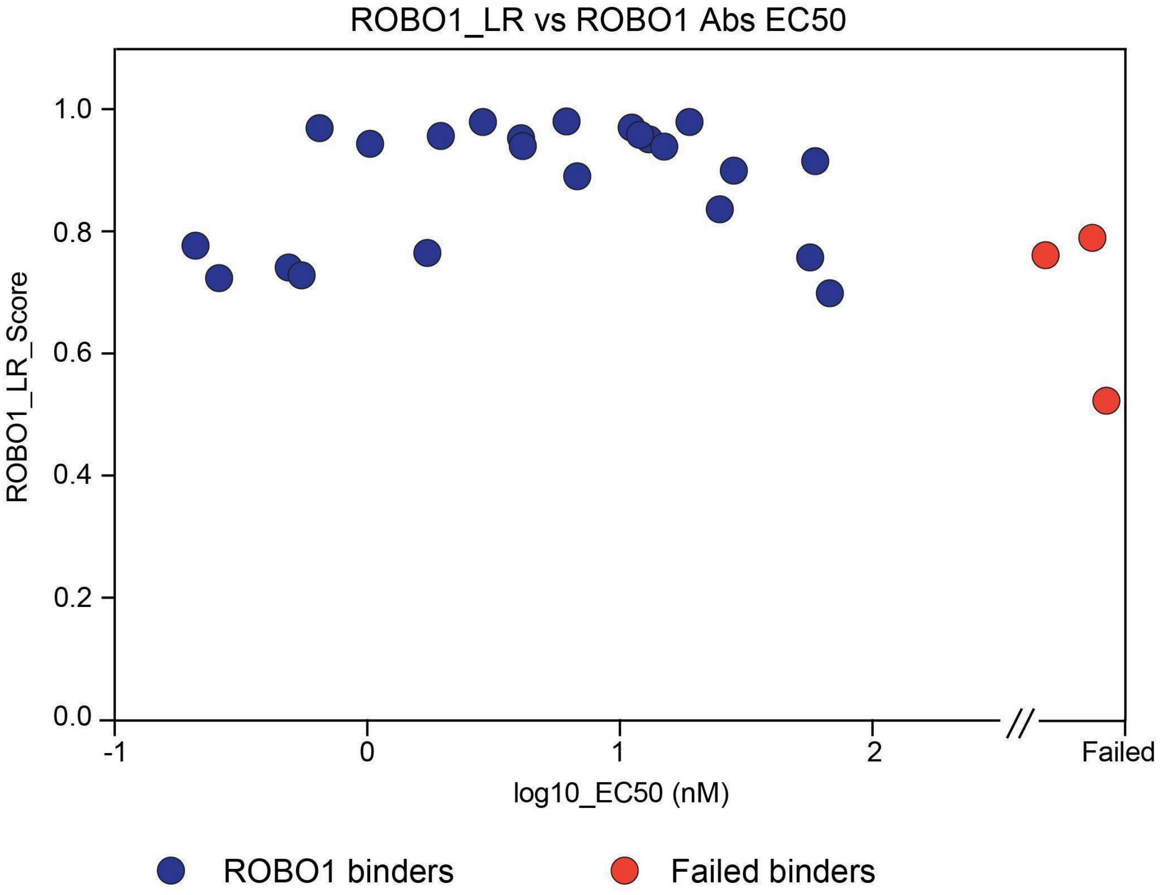


**Figure S9.** Correlation between the ROBO1_LR score (ranging from 0 to 1) and the cell display EC50 values experimentally obtained for the ROBO1 antibodies. Antibodies that failed to bind in cell display are colored red.

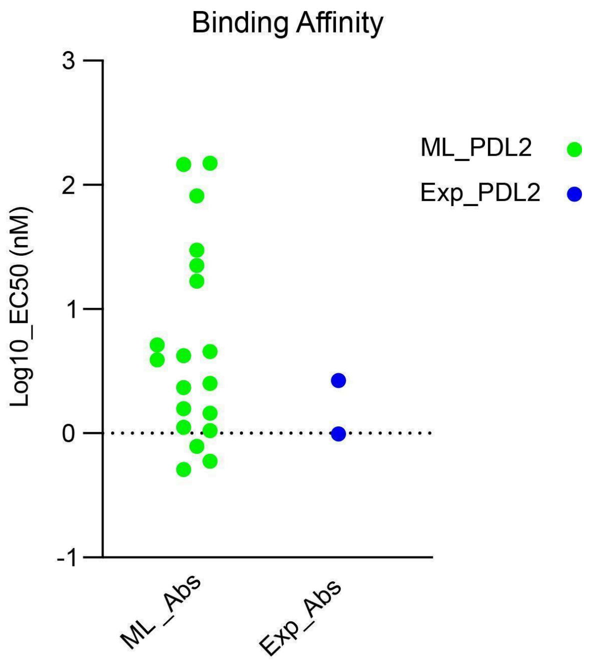


**Figure S10.** Cell display antibody titration EC50 values for the ML derived PDL2 antibodies (green) and the experimentally derived antibodies (in blue).

### References


1. Haryadi, R. *et al.* Optimization of Heavy Chain and Light Chain Signal Peptides for High Level Expression of Therapeutic Antibodies in CHO Cells. *PLOS ONE* **10**, e0116878 (2015).

2. Güler-Gane, G. *et al.* Overcoming the Refractory Expression of Secreted Recombinant Proteins in Mammalian Cells through Modification of the Signal Peptide and Adjacent Amino Acids. *PLOS ONE* **11**, e0155340 (2016).

3. Crooks, G. E., Hon, G., Chandonia, J.-M. & Brenner, S. E. WebLogo: a sequence logo generator. *Genome Res* **14**, 1188–1190 (2004).
